# Predicting the DNA binding specificity of transcription factor mutants using family-level biophysically interpretable machine learning

**DOI:** 10.1101/2024.01.24.577115

**Authors:** Shaoxun Liu, Pilar Gomez-Alcala, Christ Leemans, William J. Glassford, Lucas A.N. Melo, Xiang-Jun Lu, Richard S. Mann, Harmen J. Bussemaker

**Affiliations:** Department of Biological Sciences, Columbia University, New York, NY, USA; Department of Biochemistry and Molecular Biophysics, Columbia University, New York, NY, USA; Department of Systems Biology, Columbia University, New York, NY, USA

**Keywords:** transcription factors (TFs), DNA binding specificity, functional impact of missense mutations, basic helix-loop-helix (bHLH), homeodomain, high-throughput SELEX, protein binding microarray (PBM), biophysically interpretable machine learning

## Abstract

Sequence-specific interactions of transcription factors (TFs) with genomic DNA underlie many cellular processes. High-throughput *in vitro* binding assays coupled with machine learning have made it possible to accurately define such molecular recognition in a biophysically interpretable way for hundreds of TFs across many structural families, providing new avenues for predicting how the sequence preference of a TF is impacted by disease-associated mutations in its DNA binding domain. We developed a method based on a reference-free tetrahedral representation of variation in base preference within a given structural family that can be used to accurately predict the effect of mutations in the protein sequence of the TF. Using the basic helix-loop-helix (bHLH) and homeodomain families as test cases, our results demonstrate the feasibility of accurately predicting the shifts (ΔΔΔG/RT) in binding free energy associated with TF mutants by leveraging high-quality DNA binding models for sets of homologous wild-type TFs.

Graphical Abstract

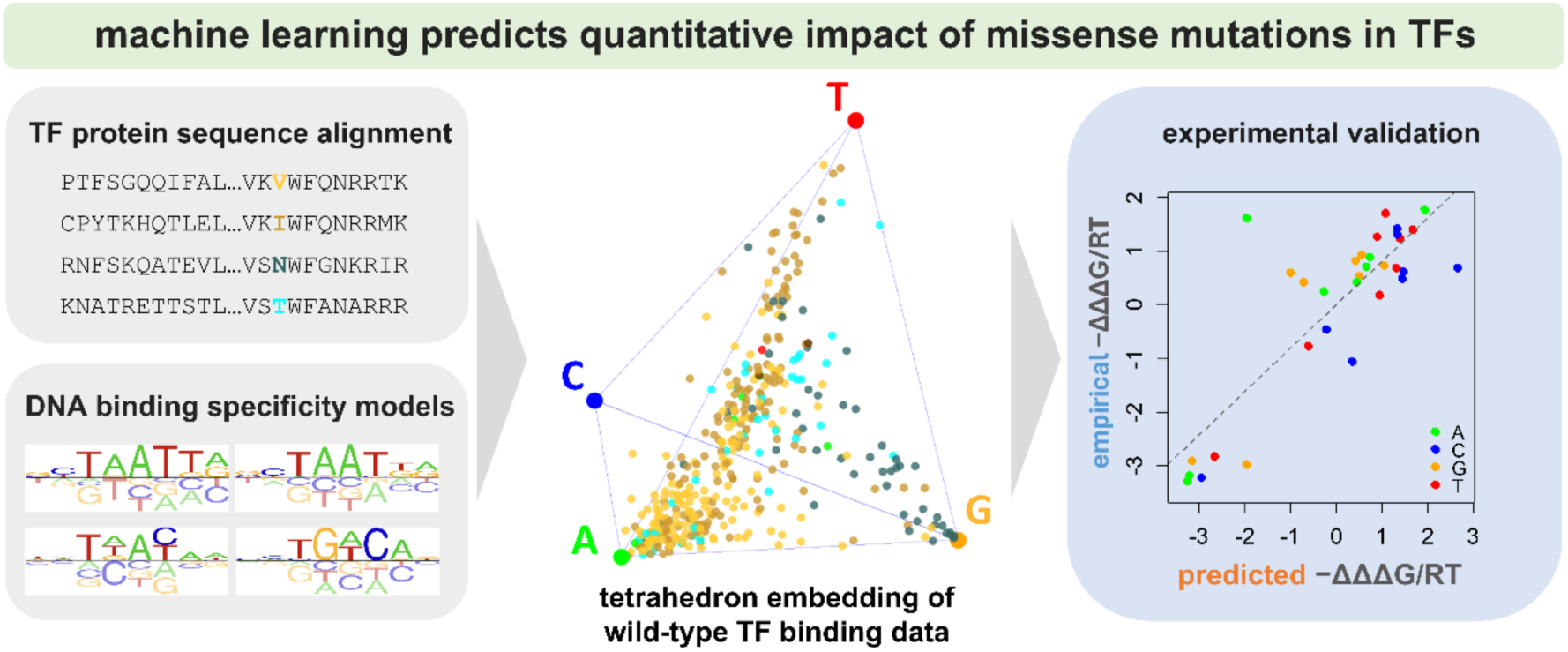

## INTRODUCTION

Gene expression regulation, mediated to a significant extent by transcription factors (TFs), is a central aspect of cellular function that is especially important during cellular differentiation and stress responses. The DNA binding domain of a TF enables it to perform its regulatory function by recognizing and binding to specific DNA sequences (1). TF binding sites typically reside either in proximal promoter regions near transcription start sites or in more remote enhancer regions. By binding to these sites, TFs regulate gene expression, frequently by recruiting transcriptional co-activators or co-repressors (2). The range of TF binding affinity – typically represented by a dissociation constant (K_D_), which equals the free TF concentration at which the DNA is bound 50% of the time – covers many orders of magnitude between optimal and non-specific binding, and even ultra-weak binding sites can be functionally relevant (3–6). It is therefore of great value to be able to predict DNA binding affinity over its full range, and how this affinity changes when TF binding sites are mutated (7).

Previous studies have provided evidence that mutations in TFs are responsible for numerous common developmental disorders. For example, anophthalmia has been linked to mutations in SOX2 (8), while autosomal dominant Rolandic epilepsy is associated with mutations in NEUROG1 (9). A single point mutation in the DNA binding domain of TFs has the potential to disrupt their DNA binding preference, leading to dysregulated developmental pathways (10, 11). In addition to disrupting binding, TF mutations can cause quantitative shifts in DNA binding affinity, which can be functionally relevant even when the preferred base remains the same (12). Detecting such subtle changes requires quantitative protein-DNA binding assays such as protein binding microarrays (PBM) (13), SELEX-seq/HT-SELEX (14–16) or chromatin immunoprecipitation coupled with deep sequencing (ChIP-seq) (17). These assays, however, are both labor-intensive and expensive: PBM and SELEX-seq involve in vitro protein purification; ChIP-seq requires manual collection of tissues and immunoprecipitation with antibodies that can vary widely in their efficacy. As a consequence, with some exceptions (18), their application has mostly been limited to wild-type TFs. The development of an accurate computational model trained on binding data for *wild-type* TFs, but capable of making predictions for *mutant* TFs, would facilitate the large-scale characterization of TF variants, as it would greatly reduce the number of variants that need to be tested experimentally.

When modeling the DNA binding specificity of an individual TF, an approximation that treats the effects of base-pair mutations at different positions within the DNA binding site as independent is often relied upon (19–22). This is equivalent to assuming additivity of binding free energies. For a given TF, the binding specificity model then takes the form of a position-specific matrix containing the free energy differences (ΔΔG in units of RT) associated with base substitutions at each DNA position. In previous work, we developed computational methods for estimating ΔΔG/RT parameters from high-throughput TF binding data with sufficient accuracy to make predictions of low-affinity binding sites functionally meaningful (6, 22, 23).

In the present study, we build on an interpretable machine learning framework called ProBound that was recently developed by our lab (22). ProBound allows us to accurately estimate the binding free energy parameters that define a TF’s base preferences using data from in vitro binding assays. While it is possible to extend this model to account for dependencies between nucleotide positions, the improvement in binding energy prediction accuracy over simple position-specific matrix representation is often only modest (6, 24), so we here restrict ourselves to simple additive free energy models. To analyze the quantitative relationship between base preference and TF protein sequence within a given TF structural family, we use an innovative reference-free tetrahedron representation of base preference. This enables us to systematically map the protein features that are the most important determinants of differences in DNA binding specificity among TF paralogs. Our method requires as inputs for model training the protein sequences of representative transcription factors within the same family along with data from suitable in vitro DNA binding assays on wild-type transcription factors. Several previous studies also analyzed variation in DNA binding specificity among wild-type TFs within specific structural families (25–29). However, recent large-scale efforts to empirically determine the impact of disease-associated missense mutations in TF protein sequences on DNA binding specificity (18, 30) have created a new opportunity to benchmark the predictive performance of such family-level predictive modeling in a biologically meaningful application. As a proof of concept, we here focus on two distinct TF families: basic helix-loop-helix (bHLH) proteins, which are widely studied in both eukaryotes and bacteria (31), and homeodomain (HD) proteins, which play major developmental roles in eukaryotes (32).

## METHODS

### Collecting HT-SELEX data for training bHLH binding models

The training set data used in this study consisted of HT-SELEX data for all TFs annotated as belonging to the bHLH family from three publications of the Taipale lab (16, 33, 34). For the data from Yin et al., we only used the unmethylated library and ignored reads corresponding to methylated DNA ligands. This yielded a total of 121 multi-round HT-SELEX datasets covering 62 distinct bHLH proteins.

### Constructing binding free energy models using ProBound

For each of the 121 HT-SELEX datasets, we ran ProBound (http://github.com/RubeGroup/ProBound and Ref. (22)) using the JSON configuration file in **Supplemental Data S5**, which let us fit a single binding energy model to data from all rounds of the SELEX assay simultaneously. We then filtered the models by model quality based on the following two heuristic criteria: (i) sufficient ability of the model to predict sequence enrichment at the level of 8-mers (R^2^ > 0.15), and (ii) a base preference pattern consistent with the E-box consensus CANNTG in the center of the binding energy model. A total of 97 models passed these criteria. An bHLH factor was assigned the binding model with the highest R^2^ value if multiple models for it passed the quality filter. Our final bHLH binding model compendium comprised 52 distinct bHLH proteins.

### Tetrahedron representation of DNA binding energies

In the position-specific affinity matrix (PSAM) representation for a given TF (35), each column *j* corresponds to a different position within the DNA binding site, each row *b* to a different base, and each element *w*_j*b*_ to the affinity relative to the preferred base *b*_0_(*j*) at position *j* (this implies that *w*_j*b*0(j)_ = 1 for the preferred base at each position). To map the four relative affinities in a column of the PSAM to a 3D position within the tetrahedral embedding space, the relative affinities are first normalized to frequencies *f*_j*b*_ = *w*_j*b*_/+∑_*b*′_ *w*_j*b*′_). In a subsequent transformation step, the transpose of the frequency matrix is multiplied by a 4×3 tetrahedral transformation matrix:

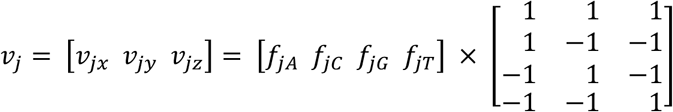

This operation maps each PSAM column containing four elements to the Cartesian coordinates (*x*, *y*, *z*) within a tetrahedron. Note that the tetrahedron has corners located at positions (1, 1, 1), (1, −1, −1), (−1, 1, −1), and (−1, −1, 1), corresponding to a strict preference for A, C, G, and T, respectively.

### Alignment of bHLH protein sequences

The cloned protein sequences of the bHLH transcription factors whose HT-SELEX date we trained on were obtained from Refs. (16, 33, 34). They were individually aligned to the hidden markov model (HMM) for PFAM entry PF00010 (https://www.ebi.ac.uk/interpro/entry/pfam/PF00010/) (36, 37). This allowed their amino-acid identities to be mapped to standardized residue positions in the range 1-53.

### MANOVA analysis

A series of MANOVA tests were performed to examine the association across all bHLH factors between amino acid identity (independent variable) at a given residue position in the protein alignment and tetrahedral position (dependent variable) reflecting the base preferences for a given position in the DNA binding model. For each combination of residue position and DNA position, we used the summary.manova() function in R, with the F-statistic computed using the Pillai-Barlett trace test. The null hypothesis is that the mean tetrahedral position is the same for all amino acids.

### bHLH replicate comparisons used as reference for predictions

Of the 52 unique bHLH proteins, 36 had two replicates available (when more than two replicates were available, we chose the two with best correlation between predicted and observed 8-mer enrichment). The ΔΔG/RT estimates from the binding model for these replicates were compared to obtain a reference for predictive performance.

### Definition of closest-paralog for predicting binding specificity

For each of the 52 bHLH proteins that was held out, the protein with the smallest Levenshtein distance in terms of protein sequence (equivalent of percentage amino-acid identity) was selected from the remaining 51 sequences. These ΔΔG/RT estimates from the binding model for this closest-paralog were then used as predicted values.

### Implementation of similarity regression prediction

Pre-trained positional weights are adopted from the original similarity regression paper for the bHLH and HD families (38). For the bHLH family, two gaps were added between the 34^th^ and 35^th^ position of the bHLH alignment to match the alignment length recorded in the similarity regression study. The difference in alignment length is likely cause by usage of different HMM models to represent the family. The most highly represented motif is selected by multiplying the AA identity profile with the pre-trained weights at each position, deriving a weighted similarity score. The motif associated with the protein sequence that has the smallest similarity score is selected as inferred motif.

### Construction of SVD-regression model

To perform SVD regression for a given DNA position, we first constructed a tetrahedral coordinate matrix *M* whose three columns correspond to the coordinates of the 3D space in which the tetrahedron is embedded, and whose rows correspond to the set of bHLH proteins analyzed. The 52×3 matrix *M* was centered by subtracting the column mean from each value. Next, we performed singular value decomposition (SVD): *M* = *UDV*^*T*^ using the base function svd() in R. The columns of 3×3 matrix *V* define a natural data-driven basis for the vector space inside the tetrahedron, while the columns of U correspond to the projection of the points inside the tetrahedron along each of these principal component (PC) directions.

For each of the three PCs (*c*=1,2,3) for a given DNA position, we separately constructed a linear regression model that predicts the position of an unseen bHLH factor from its protein sequence. Each residue position *r* in the bHLH multiple alignment corresponds to a feature that could be used as a predictor. The importance of each of the 159 combinations among 53 residue positions and 3 PCs was assessed using an ANOVA test implemented using the aov() function in R, with the amino acid identity *a* as the independent variable and the values in each column of *U* as the dependent variable. Each residue position is associated with three p-values, one for each PC. For each PC, we ranked the protein features by their ANOVA p-value, and iteratively selected for the features to include. For each principal component *c* we started by fitting a linear regression model based on the most statistically significant residue position *r*, in which the position *m*_*cf*_ along *c* of each transcription factor *f* in the centered matrix *M* was predicted to be proportional to the mean position of all TFs with the same amino acid at position *r*:

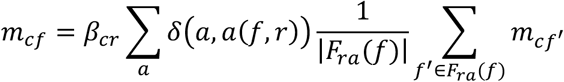

Here, *β*_*cr*_ denotes a regression coefficient, *δ*(*a*, *a*′) = 0,1 is an indicator function that is equal to 1 when *a* = *a*′ and 0 otherwise, *a*(*f*, *r*) is a lookup function that provides the amino acid identity at residue position *r* for factor *f*, and *F*_*ra*_(*f*) is the set of all TFs *except f* that have amino acid *a* at position *r*. Next, we compared with a linear model using an additional feature. As long as the p-value of the F-test was below 0.05, indicating a significant improvement in variance explained, we included the additional feature in our model, and proceed to the next feature. After the iterative feature selection step, residue positions selected for each PC will have coefficients associated with them that get estimated using linear regression, resulting in three models, one for each of the three PCs. When making predictions for an unseen TF, the mean PC value across all TFs in the training set with the same amino-acid at a given position is used.

For amino-acids unseen in the training set at a given position, the PC value is estimated as a mean across all amino-acids in the training set weighed by the reverse of the difference between the identity score and mismatch score in the BLOSUM62 matrix (39). The encoded values of each residue position are multiplied by the coefficient matrix to generate predicted PC values. These are in turn transformed into tetrahedron coordinates using a reverse SVD transformation, and mapped to PSAM using the inverse operation of the tetrahedron embedding. For cases where the resulting PSAM has one or more negative values, they are set to 0.001. Finally, the predicted PSAM is transformed to –ΔΔG/RT values by taking the natural logarithm of the relative affinities.

### Assessment of prediction confidence

A confidence score associated with each prediction for a protein sequence at a motif position was computed as follows:

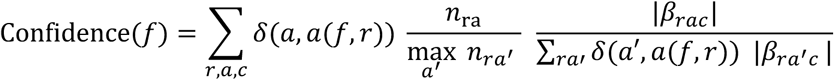

Here the sum runs over all residue positions *r*, amino acid identities *a*, and principal components *c*.

### Predicting ΔΔΔG/RT for TFs mutants using tetrahedral regularization

To predict the effect of TF point mutations on DNA binding specificity, two modifications are made to the SVD regression model. Firstly, during feature selection, protein features that represent the mutated positions are designated as key features, and iterative feature selection is not used. Secondly, instead of directly predicting the – ΔΔG/RT values for the mutant TF and subtracting the – ΔΔG/RT values for the corresponding wild-type TF, we calculated the shift between the mutant and wild-type TF in tetrahedron space and transformed the difference back to energy space to obtain ΔΔΔG/RT values, using the same transformation procedures as used for the construction of SVD-regression model above (**Figure S5**).

### Application to PBM data from the CisBP database

To train the PCA-regression model on PBM data, PWMs, TF metadata, and normalized fluorescent intensities (Z-scores) were downloaded from the CisBP database (https://cisbp.ccbr.utoronto.ca/) for both the bHLH and HD protein families. To analyze the motif models in CisBP, we directly imported all PWMs from the database and scored each motif model with the corresponding motif seed (CANNTG for bHLH, NNTDAYNN for HD) and collected the best-matching kmer motif models. For PBM entries for which z-scores were available, we constructed a binding model with PyProBound (23), using NCANNTGN and NNTDAYNN as an 8mer seed for bHLH and HD, respectively. We then searched for the sequence of the longest isoform of each unique protein from Uniprot (40), and aligned it to the HMM of the bHLH and HD families from PFAM (37). A detailed description of the analyses with reproducible R scripts is available at https://github.com/BussemakerLab/FamilyCode.

### Wild type and mutant HES2 and ASCL2 protein expression

ASCL2 and HES DNA binding domain (DBD) sequences were expressed based on the cloned sequences described in Yin et al. (34), tagged with an N terminal 6xHis tag, and expressed and purified according to the protocol described in Slattery et al. (41). To create mutant proteins, site-directed PCR was performed using primers containing mutations at the 5th and 13th residue positions. For HES2, the wild type and mutant sequences were cloned into the pQE30 vector (Qiagen). ASCL2 wild type and mutants were tagged with mScarlet to enhance solubility (42), and were cloned into the pET11 vector (Novagen). The full mScarlet sequence is as follows:

VSKGEAVIKEFMRFKVHMEGSMNGHEFEIEGEGEGRPYEGTQTAKLKVTKGGPLPFSWDILSPQ FMYGSRAFIKHPADIPDYYKQSFPEGFKWERVMNFEDGGAVTVTQDTSLEDGTLIYKVKLRGTN FPPDGPVMQKKTMGWEASTERLYPEDGVLKGDIKMALRLKDGGRYLADFKTTYKAKKPVQMPGA YNVDRKLDITSHNEDYTVVEQYERSEGRHSTGGMDELY

All protein samples are expressed in BL21 E. coli cells. The BL21 cell are grown in LB plus 100μg/ml carbenicillin for 2.5 hours before IPTG induction and continues to express proteins for ∼4.5 hours at 37 ℃. The cells are then collected by centrifugation and resuspended in 8ml of lysis buffer (50mM Tris pH 7.5, 600mM NaCl, 40mM Imidazole). The lysate then went through sonication for 5 rounds of 30 seconds with 1-minute intervals. The lysate was then spun down and proteins in the supernatant are purified with affinity purification using Cobalt-Talon beads (Clontech) using a low imidazole wash buffer (50mM Tris pH 7.5, 600mM NaCl, 20mM Imidazole), and collected with a high imidazole elution buffer (50mM Tris pH 7.5, 600mM NaCl, 300mM Imidazole). Protein samples are dialyzed in dialysis buffer (20mM HEPES pH 7.9, 200mM NaCl, 10% Glycerol, 2mM MgCl_2_) overnight before changing dialysis buffer and another 2-hour dialysis. All procedures above are performed at 4℃.

The final concentration of HES2 obtained was 17±3μM for wild type and R5K mutant and 70±20μM for R13V and double mutant; the final concentrations obtained were 14±1μM for ASCL2 wild type and mutants.

### SELEX assays

SELEX experiment were performed following the protocol outlined in Riley*, et al.* (14) and a SELEX library with a 16-mer randomized region whose full sequence is as follows:

GTTCAGAGTTCTACAGTC-CGACCTAA-16N-TTAGG-ACTCGGACCTGGACTAGG

The libraries were double strand DNA annealed and extended with Klenow polymerase. The SELEX annealing primers are as follows:

SELEX-L - GTTCAGAGTTCTACAGTCCGA SELEX-R – CCTAGTCCAGGTCCGAGT

To be specific, the initial mixture was prepared in a DNA LoBind Safe lock tube (Eppendorf 022431021) by combining 10μL of 10x STE buffer (100mM Tris pH8.0, 10mM EDTA pH8.0, 1M NaCl), 10μL of 100 μM SELEX library oligo, 20μL of 100μM annealing primer (sequence), and 60μL of water. The tube was then placed in a 1-liter beaker containing 800mL of water and boiled for 10 minutes. After boiling, the tube was left in the hot water until the water cooled to room temperature naturally. Next, 100μL of the annealed DNA from the first step was taken and mixed with 25μL of 10x Klenow buffer, 20μL of 10mM dNTP, 80μL of water, and 20μL of Klenow fragment (NEB M0210L). The mixture was incubated at room temperature for 30 minutes. To stop the reaction, 10μL of 0.5M EDTA at pH 8.0 was added. Following this, 25μL of 3M sodium acetate at pH 5.2 was added, and the DNA was purified using a Qiagen column (Qiagen 28106) according to the regular PCR purification protocol.

With the SELEX libraries ready, an EMSA gel was prepared to separate the bound DNA fragments from the unbound. To prepare a gel for EMSA, 3.5ml of 5× TBE, 3.1ml of 30% acrylamide/bis-acrylamide solution (37.5:1), 2.3ml of 40% acrylamide solution, and 1.1ml of 80% glycerol were added to 24.25ml of water. The mixture was mixed well, avoiding the generation of bubbles. Then, 262.5mL of 10% ammonium persulfate and 17.5mL of TEMED were added to catalyze polymerization. The solution was briefly mixed, poured into a mold to form a gel, and allowed to solidify for 1 hour. After solidification, the wells of the gel were flushed with 0.5× TBE to remove unpolymerized acrylamide, and the gel was pre-run for 10 minutes at 150 V in 0.5× TBE buffer. The gel was run in a cold room at 4°C.

Fluorescent EMSA assays were performed alongside the SELEX reactions to define the position of the bound band in the gel. The high-affinity probes for HES2 and ASCL2 binding, respectively, were as follows:

HES2 CTCTCCTCCGTCAA**<u>CACGTG</u>**TTGAGCAGCGCAGTCGTATGCCGTCTTCTGCTTG

ASCL2 CTCTCCTCCGTCAA**<u>CAGCTG</u>**TTGAGCAGCGCAGTCGTATGCCGTCTTCTGCTTG

For the Fluorescent EMSA reactions, 30mL reactions were set up as follows: 9μL of labeled control probe at 166.67nM final concentration diluted with water, 15μL of ASCL2 wild type or HES2 wild type protein at 300nM final concentration diluted with dialysis buffer, and 6mL of 5× binding buffer (50mM Tris pH 7.5, 250mM NaCl, 5mM MgCl_2_, 20 % glycerol, 2.5mM DTT, 2.5mM EDTA pH 8, 250μg/ml polydI-dC, 1mg/ml BSA.).

For the SELEX binding reaction, 30mL reactions were set up as follows: 9μL of SELEX library diluted with water at final concentration of 200nM, 15μL of HES2 or ASCL2 wild type or mutant protein diluted with dialysis buffer at final concentration of 133.33nM, and 6 microliters of 5× binding buffer.

Both control and SELEX binding reactions were assembled and incubated for 30 minutes at room temperature. The EMSA gel was pre-running during the binding reactions for the last 10 minutes. After the incubation, the pre-running of the gel was stopped, samples were loaded into the lanes, and the gel was run for approximately 100 minutes at 150 V (4°C).

After the run stopped, the bound bands were cut out (**Figure S12**), purified, PCR amplified, and sequenced with NextSeq 500 according to protocol from Kribelbauer *et al.* (43).

### SELEX data analysis

ProBound (22) was applied for all 8 protein samples to construct binding free energy models using the JSON configuration file in **Supplemental Data S5**, resulting in symmetrical motif matrices with a CANNTG E-box core.

### Wild type and mutant ASCL2 EMSA experiment

EMSA assays were performed using ASCL2 wild type and mutant protein and 32P-labeled DNA probes. Protein samples were prepared using the same procedure as above. The DNA probes used, however, were different to avoid affinity towards the flanking regions:

CACGTG probe:

TAGCCAATAACTTCGTCCCT**<u>CACGTG</u>**CATATAAGGAAGATCTAACCACCAATTTGG

CAGCTG probe:

TAGCCAATAACTTCGTCCCT**<u>CAGCTG</u>**CATATAAGGAAGATCTAACCACCAATTTGG

Each probe was synthesized with P32 labeled SRI sequence (CCAAATTGGTGGTTAGATCTTCC, reverse complementary to the end of the probe sequence) through annealing and Klenow reactions.

50nM of probe was combined with 500nM, 1μM, and 2μM of protein, respectively, in each binding reaction, and the same reagents as used in Feng et. al., (44). The mixture was loaded into a polyacrylamide gel after 30 minutes of reaction time. An EMSA gel was run for 100 minutes at 150 V (4°C), and dried and imaged using a phosphor-imaging plate and Typhoon gel imager.

ΔΔG/RT measurements of a probe changing from CACGTG to CAGCTG of each ASCL2 wild-type or mutant sample is calculated by measuring the band intensities of the bound and unbound band using ImageJ and fitting the following equation:

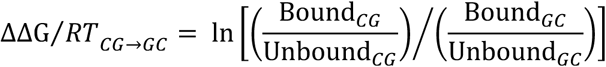

Here *Bound* and *Unbound* represent the measured band intensities of the corresponding lanes.

### Application to homeodomain data

Binding affinity models built from HT-SELEX data for HD factors using ProBound were obtained from MotifCentral.org (22). When using PyProBound to infer binding models from PBM data, as a pragmatic way to impose a consistent “binding frame” on the models, we aligned each of the resulting HD models to the NNTDAYNN 8-mer IUPAC consensus motif for HDs (here D matches A, C, or T; Y matches the pyrimidines C or T only, and N matches any base) by optimizing a motif matching score (MMS) defined as follows:

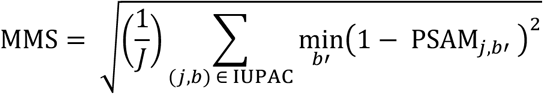

Here *L* is the length of the motif seed consisting of a 4x*L* matrix representing the IUPAC identification of the motif. The 4 rows of the matrix represent the 4 bases, and each column represents a motif position. Each position in the matrix is set to unity if the IUPAC symbol covers the given base, and zero otherwise. The protein sequences of the HD proteins were retrieved as the recorded cloned sequences, and aligned to the HMM model for PF00046 (https://www.ebi.ac.uk/interpro/entry/pfam/PF00046/) (36, 37) using the same procedure as for the bHLH family. Note that with this definition of residue position, the traditional Arg51 based on the Engrailed sequence (45) is now Arg50 (**Figure 5A, Figure S9**).

### Analysis of PBM data for mutant homeodomains using PyProBound

We obtained protein-binding microarray (PBM) data for 92 single-residue mutants of human homeodomain (HD) proteins and their corresponding wild-type HDs from Ref. (30). Raw intensity data were processed using PyProBound to generate ΔΔG/RT models. The quality of each model was assessed by calculating the coefficient of determination (R²) between predicted and empirical probe intensities (22, 46, 47). Of the 92 mutants reported, 89 had at least two replicates. For each mutant, the replicate with the highest R² was designated as the replicate 1, and the replicate with the second-highest R² as replicate 2. Empirical ΔΔΔG/RT values were determined for each mutant replicate by comparing its ΔΔG/RT model to its wild-type counterpart using the tetrahedron regularization. The R^2^ and RMSD between the ΔΔΔG/RT values for the respective replicates served as a measure of empirical reproducibility.

### Using FamilyCode to make ΔΔΔG/RT predictions for homeodomains

We used a methodology similar to how we evaluated the HLH-1 mutants. A FamilyCode model was trained on a compendium of ΔΔG/RT binding specificity models for wild-type HDs derived from a mix of SELEX and PBM data, and used to predict ΔΔΔG/RT values quantifying base preference shifts for each of the 92 mutant HDs from Ref. (30). Empirical ΔΔΔG/RT values were computed by using the tetrahedron representation to compare the ΔΔG/RT model derived from the PBM data (30) for each mutant HD with its wild-type counterpart. Prediction accuracy was evaluated by computing the R² and RMSD between predicted and empirical ΔΔΔG/RT values across all eight motif positions.

### Comparison with rCLAMPS

The rCLAMPS tool was implemented following instructions from its GitHub repository (48). Full-length sequences of each HD protein from the study were retrieved from UniProt (40), selecting the longest isoform, and point mutations were introduced as specified. A FASTA file containing the mutant sequences was input into the pretrained rCLAMPS model, yielding 92 PWM motif models. These models were used to calculate ΔΔΔG/RT values and evaluated in the same manner as our FamilyCode models with the slight modification of using 6 nucleotide positions instead of 8, as rCLAMPS only predicts binding specificity to the TDAYNN 6mer motif. To assess the difference between the prediction accuracy of the FamilyCode model and the rCLAMPS model, we performed a paired Wilcoxon signed-rank test on the 92 R^2^ values.

### Comparison with DeepPBS

Unlike FamilyCode and rCLAMPS, the DeepPBS method (49) relies on structural information to predict DNA binding specificity for unseen TFs. We obtained full-length sequences of mutant HD proteins from UniProt, selected the longest isoform, and used the same DNA sequence as an initial guess for HD proteins (GCGTGTAAATGAATTACATGT) as in Ref. (49). These sequences were submitted to the AlphaFold3 server (50) to predict the structures of the protein-DNA complexes. Model_0 from each prediction was then submitted to the DeepPBS server with the “both readout” option to obtain PWM predictions. The predicted motifs were aligned to the NNTDAYNN 8-mer core using the same method we used for the HD training data alignment. The resulting motif models were used to calculate ΔΔΔG/RT values using the tetrahedron representation.

## RESULTS

### A collection of SELEX-derived DNA binding specificity models for wild-type bHLH factors

bHLH proteins function as homo- and/or heterodimers that prefer to bind to sequences whose core matches the reverse-complement symmetric consensus pattern CA<u>NN</u>TG (with “N” denoting any of the four nucleobases) called the enhancer box or E-box (51). We collected in vitro DNA binding datasets for 147 human and mouse DNA binding domains from the bHLH family (16, 33, 34) and used ProBound (22) to obtain biophysically interpretable and accurate binding energy models. Because bHLH homodimers are expected to have palindromic binding preferences (meaning that the binding free energy differences associated with base-pair substitutions are subject to reverse complement symmetry, but not that the binding sites themselves need to be palindromic), we configured ProBound to impose reverse-complement symmetry on its binding free energy parameters (22). This yielded a binding model that matches the symmetrical CA<u>NN</u>TG consensus E-box for 96 out of the 147 experiments analyzed, covering 54 distinct bHLH proteins. Further filtering for high intrinsic model quality resulted in a compendium of DNA recognition models for 52 bHLH factors (**Figure 1** and **Supplemental Data S1**). For each TF included in this training set, the cloned sequence for each protein covers the 53 amino acid bHLH HMMER profile, with a consensus Glu residue at position 9 (31). With only a few exceptions, the preferred DNA binding site contains one of three E-box variants (CA<u>CG</u>TG, CA<u>GC</u>TG, or CA<u>TA</u>TG). Protein sequences were aligned using the HMMER (52) profile of the bHLH family; alignment of the DNA binding specificity models was trivial thanks to their imposed palindromic symmetry. See **Supplemental Methods** for details.

**Figure 1:**
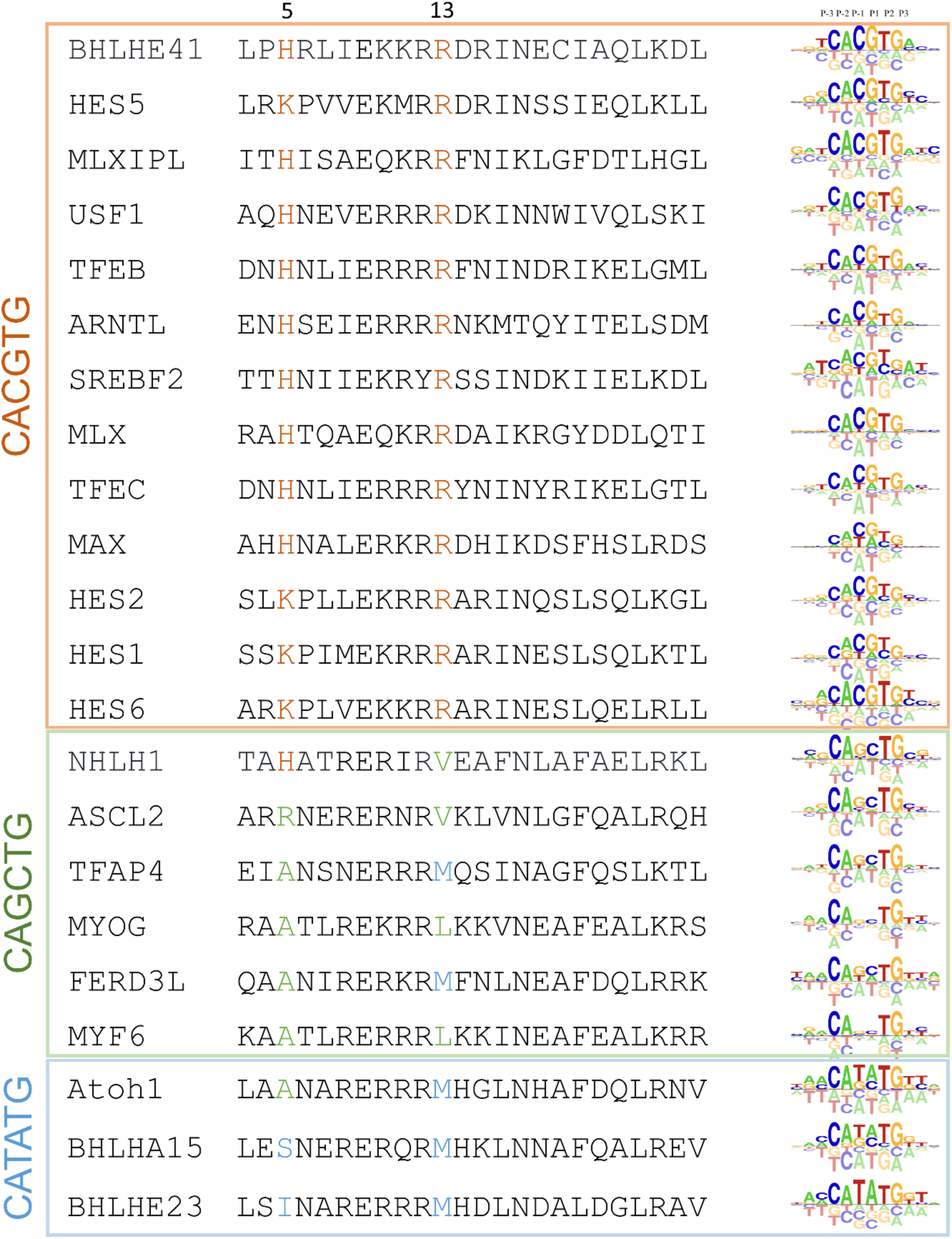
Variation in DNA binding preference within the helix-loop helix (bHLH) family of transcription factors. Aligned protein sequences and DNA binding energy logos for a representative set of bHLH factors, grouped by preferred E-box core.

### A reference-free and robust tetrahedron representation of DNA base preference

The DNA binding specificity of each TF in **Figure 1** is defined in terms of a matrix of binding free energy differences ΔΔG associated with base substitutions at each position in the DNA binding site. Setting ΔΔG=0 for the preferred base leaves three independent ΔΔG parameters for each position in the DNA binding site. The corresponding relative affinity parameters are equal to exp(–ΔΔG/RT) and together constitute a position specific affinity matrix (PSAM) (35). When the goal is to explain variation in ΔΔG/RT in response to variation in the amino acid sequence across set of TFs from the same family, the preferred base at a given position will not necessarily be the same for each TF. Moreover, for bases that are strongly disfavored at a given position, the corresponding ΔΔG/RT values as estimated from the binding data can have large fluctuations that are not biophysically meaningful as they all correspond to a relative affinity near zero. These technical issues are addressed by mapping the ΔΔG/RT values to a three-dimensional position within a tetrahedron whose vertices correspond to the four bases (**Figure 2A** and **Supplemental Data S2**; see **Supplemental Methods** for details). Proximity of a particular TF to one of the vertices reflects a preference for the corresponding base.

**Figure 2:**
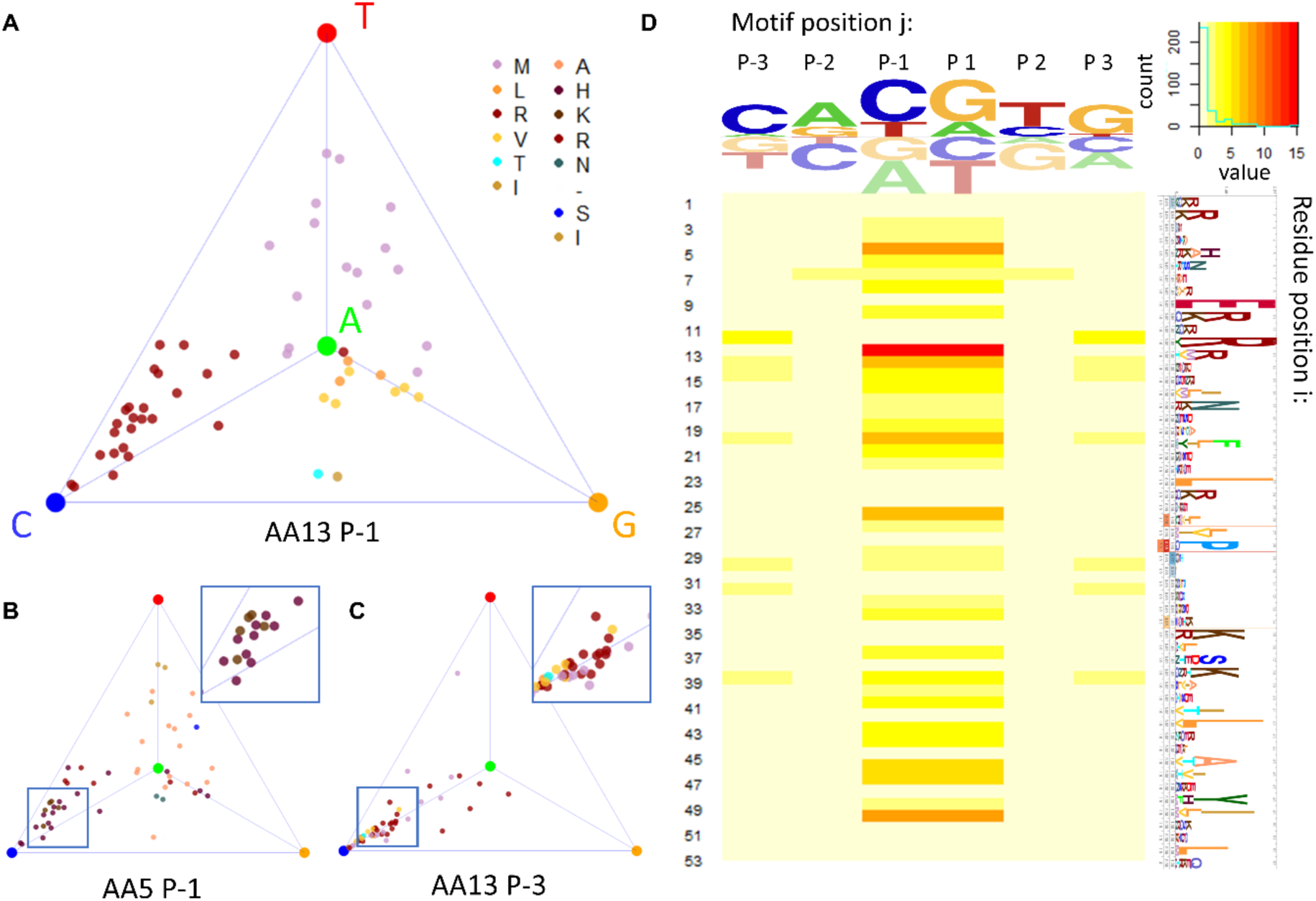
Family level analysis uncovers protein sequence determinants of DNA binding specificity. (**A**-**C**) Tetrahedron representation of base preference at DNA position –1 (panel A, B) or –3 (panel C). Each point represents a different bHLH factor, and is colored according to the amino acid identity at protein position 13 (panel A, C) or 5 (panel B). (**D**) Heatmap showing the p-value from a MANOVA test between amino-acid identity and position within the tetrahedron.

### Mapping the protein sequence determinants of DNA binding specificity for bHLH factors

As a first step towards performing family-level analysis of how the DNA binding specificity of bHLH TFs is determined by their protein sequence, we focused on DNA position –1 in the PSAM for each TF and labeled/colored each corresponding point in the tetrahedron according to the amino-acid identity at residue position 13 in the protein sequence alignment (**Figure 2A**). Visual inspection suggests that there is a trend for TFs with an arginine at protein position 13 to prefer binding to DNA sequences with a cytosine at nucleotide position –1, while a methionine at the same protein position confers a preference for thymine. A similar plot can be generated for other combinations of DNA binding site position and protein alignment position. When a different residue position is chosen in the alignment of TF protein sequences, the coloring of the points changes (**Figure 2B**); when a different position within the DNA binding site is chosen, the position of each point changes (**Figure 2C**).

To systematically dissect the relationships between amino-acid identity at residue positions in the bHLH protein sequence alignment and base preference at nucleotide positions within the DNA binding site in a way that that also assesses statistical significance, we performed a series of multidimensional analysis of variance (MANOVA) tests. Each MANOVA considers the positions of points within the tetrahedron that represented the binding affinities at a specific motif position. The three-dimensional tetrahedral coordinate plays the role of dependent variable in these tests, while the (categorical) independent variable is the amino-acid identity at a given position in the TF protein alignment. For each combination of protein and DNA position, the MANOVA yields a p-value that quantifies the statistical significance of the functional association. Because the binding model is reverse-complement symmetrical, the p-values for nucleotide positions –1, –2, and –3 are the same as +1, +2, and +3, respectively (**Figure 2D**).

We found that of the 53 residue positions in the bHLH protein alignment, 9 are significantly associated with base preference at nucleotide position –1/+1 (p < 0.001 after Bonferroni correction (53)). Visualizing the p-values from the MANOVA test along the structure of PHO4 (**Figure S1A**) shows that, as expected, the residues in direct contact with the DNA major groove (e.g., positions 5, 13, 14, and 15) tend to have the most significant associations. Base preference at nucleotide position –1/+1 is greatly influenced by amino acid identity at residue position 13 (**Figure S1B**), consistent with the fact that in PHO4, the sidechain of Arg13 directly interacts with the base-pairs at position –1 and +1 through hydrogen bonds that are known to be crucial for standard E-box preference (54, 55). Available structures of bHLH-DNA complexes with other amino acids at residue position 13 also provide a mechanistic rationale for the observed difference in base preference (**Figure S1C-E**).

Our functional analysis suggests that amino-acid variation at residue positions not in direct contact with DNA can also affect base preference. We analyzed structural data to find plausible mechanistic explanations for these cases. The residue at position 20 mediates a contact at the homodimerization interface. Having Phe or Met here promotes a preference for G and T at DNA position –1, possibly due to the larger side-chain size, while the smaller side chains of Ile and Leu confer a preference for C (**Figure S2A, B**). Structural data of protein-DNA complexes for PHO4 (PDB ID 1A0A) and Myod1 (PDB ID 1MDY) shows that for Phe20 and Leu20, the dimerization angle at the DNA binding residues is different between CACGTG and CAGCTG, with the distance between the C-alphas of Glu9 equal to 19.1 Å for CACGTG and 19.9 Å for CAGCTG (54, 56). At position 50, also distant from the DNA, the positively charged Lys and Arg, as well as Gln, are associated with a preference for C at DNA position –1, likely due to a propensity to form an electrostatic and hydrogen bond with Glu, Gln, or Asp at residue position 22. The tighter interaction between the two monomers promotes narrower docking onto DNA, which may facilitate C recognition (**Figure 1**). Among the non-DNA contacting residue positions, position 20 and 50 have not been previously associated with bHLH binding specificity, while positions 26, 46, and 47 still lack mechanistic explanations despite showing statistical significance in our analysis.

### A tetrahedron-based strategy for predicting DNA binding specificity from TF protein sequence

Beyond identifying positional correlations across the protein-DNA interface, the tetrahedron representation provides a natural starting point for predicting the effect of protein mutations on DNA binding specificity in a quantitative manner. For example, to estimate the effect of a point mutation from Arg to Val at residue position 13 in the bHLH domain on base preference at nucleotide position –1/+1, we can compute the centroid of all TFs with Arg13 or Val13, respectively (**Figure 3A, B**), and then use the vector connecting the two centroids as a predictor of the change in binding specificity upon R13V mutation of any bHLH protein that contains an Arg at position 13.

**Figure 3:**
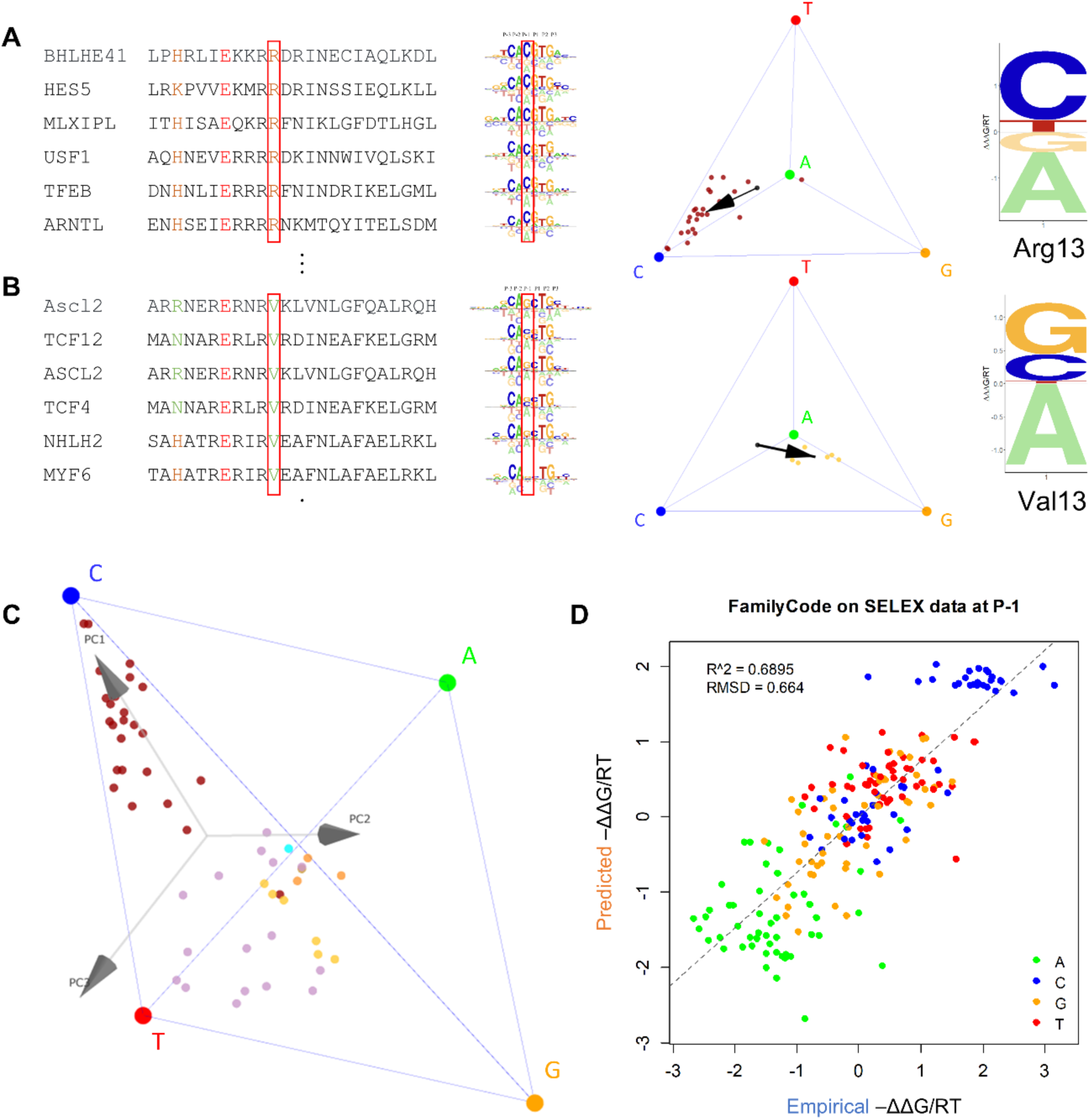
Predicting shifts in DNA binding preference associated with specific protein sequence features, and using PCA-regression to predict the binding specificity of mutated bHLH proteins. (**A**-**B**) Protein alignment and binding motif of select bHLH proteins with either Arg (panel A) or Val (panel B) at residue position 13 (highlighted by red box). The energy logos and tetrahedron plots and the right show the base preference at DNA position –1 (red box). Arrows in the tetrahedrons indicate the shift relative to the overall centroid associated with the Arg13 and Val13 subset, which can again be represented by an energy logo (far right). (**C**) Principal component analysis of the set of tetrahedral coordinates defines a natural coordinate frame for each DNA position. (**D**) FamilyCode prediction performance on individually held-out wild-type bHLH proteins.

To generalize this approach so that predictions can be made for any TF in the bHLH family, two issues need to be addressed. First, we would like to define a tetrahedral coordinate frame in a data-driven way, so that it optimally reflects the variation in chemical features that drives the interactions between amino-acid side chains and the DNA ligand. To address this, we first perform principal component analysis (PCA) on the cloud of points within the tetrahedron (see **Supplemental Methods** for details). Second, we need to account for the fact that evolutionary selection has created dependencies between amino-acid identities at distinct residue positions, which can confound the analysis when using multiple protein features simultaneously as a predictor of base preference. For instance, it would not be accurate to simply add up the individual effects of amino-acid substitutions at two residue positions that are in perfect linkage disequilibrium with each other. To address this, we perform iterative feature selection using an F-test p-value threshold for each principal component (see **Supplemental Methods** for details).

Each of the three principal components (PC) defines a natural direction within the tetrahedron onto which the variation in base preference can be projected (**Figure 3C**). Grouping TFs by amino-acid identity at a given residue position again reveals interpretable patterns (**Supplemental Data S3**). For instance, TFs containing protein feature Arg13 and Val13, respectively, are on opposite ends along PC1 (**Figure 3C)**. By performing a (one-dimensional) ANOVA for each individual principal component, functionally informative residue positions can be mapped (**Supplemental Data S4**). The difference between the mean across all TFs and the mean of the subset that matches a specific protein sequence feature such as Arg13 reflects the quantitative shift in base preference associated with that feature. Whenever a particular amino acid in the predicted TF is not present in the training data, we estimate its centroid as the mean of the centroids of all other amino acids at the same position, weighted by amino-acid substitution distance according to BLOSUM62 (39).

### Cross-validation of specificity predictions on held-out wild-type bHLH factors

As an initial assessment of this prediction scheme, which we refer to as FamilyCode, we performed leave-one-out cross-validation across the bHLH family, in which we built a PCA-regression model from all TFs except one, and then used this model to predict the tetrahedral position of the held-out TF. The prediction accuracy was 0.69 in terms of coefficient of determination (R^2^) and 0.66 in terms of root-mean-squared deviation (RMSD) of ΔΔG/RT (**Figure 3D**). For reference, we also computed these metrics of similarity between technical replicates across a subset of 20 bHLH factors for which we had access to two HT-SELEX replicates (R^2^ = 0.72, RMSD = 0.54, **Figure S3A**). We compared with a previously proposed strategy that simply takes the DNA binding specificity of the closest-paralog as the prediction (57), as well as with a refinement of this approach called similarity regression (38). Our PCA-regression model outperforms both schemes in predicting for the DNA binding energies at the –1/+1 position by a significant margin (closest-paralog: R^2^ = 0.63, RMSD = 0.75, p-value < 10^−24^, bootstrap t-test; similarity regression using pretrained weights: R^2^ = 0.58, RMSD = 0.79, p-value < 10^−33^, bootstrap t-test). See **Figure S3B**, **C** for details. Note that unlike our feature-based approach, neither of the alternative prediction methods provides detailed information about dependencies between specific DNA positions and specific TF protein positions.

When inspecting the cross-validation runs, we noticed that lower predictive performance tends to be associated with protein features that are either not covered in the training data (so that approximations need to be used that could have lower accuracy) or only sparsely covered (which causes their associated effect on base preference to be less precisely estimated from the training data). To address this, we compute a prediction confidence score as a weighted mean of the proportion of representation across all residue positions. The score ranges from 0 to 1, with 0 representing the case when none of the features are represented in the training set, and 1 representing that when each feature is maximally represented (see **Supplemental Methods** for details). We found that it is highly correlated with the actual prediction accuracy (R^2^ = 0.86; see **Figure S4**).

### Experimental validation of specificity shift predictions for mutant bHLH factors

Since high-throughput in vitro binding data are available for most human wild-type TFs (16, 33, 34), a more relevant application of FamilyCode is to predict whether and how DNA binding specificity is modulated by a particular mutation in the TF protein sequence. The number of natural TF variants in the human population is far too large for an experimental approach to be practical in that case.

We wished to perform experimental validation of FamilyCode using two different bHLH proteins: HES2, which energetically prefers the standard E-box CA<u>CG</u>TG (cytosine at position –1), and ASCL2, which prefers the E-box variant CA<u>GC</u>TG. Among other differences, wild-type HES2 contains Lys5 and Arg13 whereas wild-type ASCL2 has Arg5 and Val13 (**Figure 4A**). We first performed SELEX-seq on wild-type HES2 and ASCL2 (see **Supplemental Methods**) and used *ProBound* to infer binding models (**Figure 4C**). Comparing with models inferred from existing HT-SELEX data for the same TFs (**Figure 4B**), we found that while the ΔΔG/RT coefficients for ASCL2 were very similar (RMSD = 0.32), those for HES2 were less reproducible (RMSD = 0.61). This pointed to ASCL2 being the more suitable TF for assessing the accuracy of our predictions.

**Figure 4:**
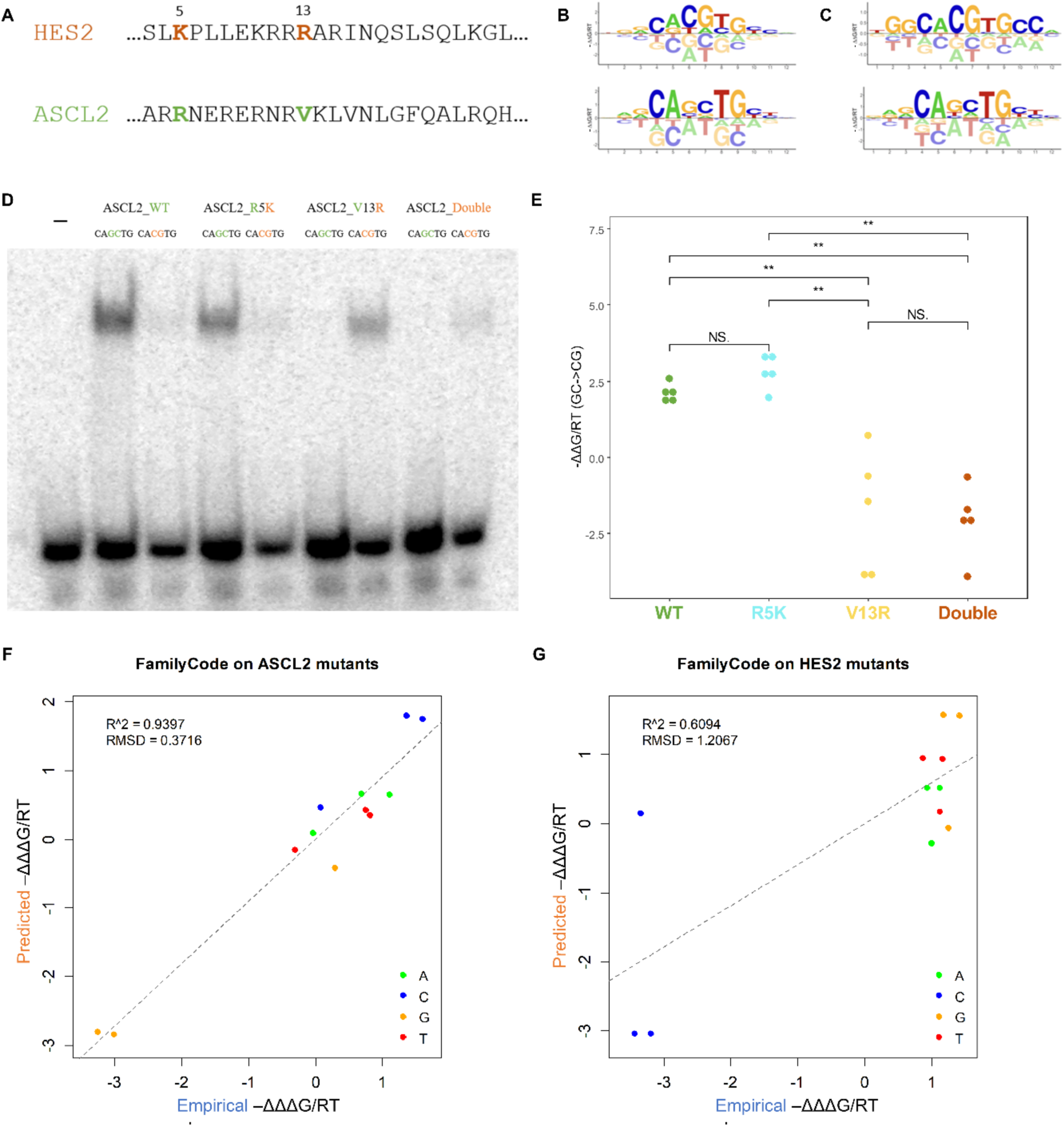
FamilyCode predictions for mutant TFs. (**A**) Sequences of wild-type human bHLH factors HES2 and ASCL2, with mutated positions 5 and 13 indicated. (**B**) Binding energy logos inferred from HT-SELEX data for wild-type HES2 and ASCL2 using ProBound. (**C**) Binding energy logos inferred from SELEX experiments for wild-type HES2 and ASCL2 using ProBound. (**D**) EMSA assays showing the binding preference between DNA probes CACGTG and CAGCTG, respectively, for both wild-type and mutant ASCL2 protein. (**E**) Statistical analysis of the changes in EMSA gel band intensity in panel D. (**F**) Predicted versus experimental values of ΔΔΔG for HES2 mutants K5R, R13V, and K5R+R13V. (**G**) Same for ASCL2 mutants R5K, V13R, and R5K+V13R.

We first used FamilyCode to predict the base preferences at position –1/+1 for ASCL2 proteins carrying the mutations R5K and/or V13R. In making these predictions, we treated each mutant as an unseen protein and predicted its binding specificity (in terms of ΔΔG/RT) the same way as for held-out wild-type TFs above. To experimentally validate these predictions, we performed electrophoretic mobility shift assays (EMSAs) with wild-type and mutant ASCL2 protein (see **Supplemental Methods**). Consistent with our predictions, these assays showed that the V13R single mutant and the R5K+V13R double mutant of ASCL2 had higher affinity for the canonical E-box (CACGTG) than the alternative E-box (CAGCTG) preferred by wild-type ASCL2 and its R5K mutant (**Figure 4D** and **4E**).

When considering a missense mutation in the protein sequence of a TF, the effect on base preference at a given DNA position is defined by a set of ΔΔΔG/RT values that quantify the *shift* in ΔΔG/RT values from the reference TF to the mutant TF (**Figure S5**). Here again, we can control the undue influence of large fluctuations in the estimated value of unfavorable bases by using the tetrahedral representation to estimate ΔΔΔG/RT, rather than directly subtracting the ΔΔG/RT values for the mutant and reference from each other (see **Supplemental Methods**).

We retrained our FamilyCode model on all available wild-type bHLH data while only fitting amino-acid substitution coefficients for the 5^th^ and 13^th^ protein positions. Predictions for the HES2 and ASCL2 mutants were made in terms of ΔΔΔG/RT values at DNA positions –1 and +1. We performed additional SELEX-seq assays for the four single mutants HES2 (K5R or R13V) and ASCL2 (R5K or V13R) and two double mutants HES2 (K5R+R13V) and ASCL2 (R5K+V13R), and applied *ProBound* to obtain a binding model for each variant. Empirical values for ΔΔΔG/RT were derived using a procedure based on the tetrahedron embedding, which is less sensitive to the larger error in binding free energy value for disfavored bases (see **Supplemental Methods**). High prediction accuracy for the variant effects was observed for ASCL2 mutants (**Figure 4F**; R^2^ = 0.94 and RMSD = 0.37). The lower prediction accuracy for HES2 mutants (**Figure 4G**; R^2^ = 0.61 and RMSD = 1.21) is consistent with the lower reproducibility of the wild-type HES2 model noted above.

### Extension of our approach to wild-type and mutant bHLH data from the PBM platform

Our family-level prediction model for the bHLH family so far relied on DNA binding specificity models for individual TFs inferred from count-based high-throughput SELEX data. To leverage the large existing body of fluorescence-based protein binding microarray (PBM) data (13), we can also directly fit ΔΔG/RT parameters to raw PBM data via a non-linear biophysical model, as originally implemented by the MatrixREDUCE (21) and FeatureREDUCE (47) algorithms. Specifically, we extended the functionality of our recent *PyProBound* (46) implementation of *ProBound* (23) from a Poisson-based to a Gaussian-based maximum-likelihood model to allow direct analysis of PBM probe fluorescent intensities (see **Supplemental Methods**). This allowed us to repeat the family-level analysis for the bHLH family using data from a distinct experimental platform and for a different set of individual TFs (most of the SELEX-based bHLH models in MotifCentral are for human TFs; most of the PBM data for bHLH factors available through CisBP are for *C. elegans* factors).

We obtained normalized PBM data for 88 bHLH domains from CisBP (57–59), constructed sequence-to-affinity models for each TF using PyProBound while imposing palindromic symmetry, and aligned protein sequences to the PFAM model of bHLH. For 80 of these bHLH domains, at least two replicate datasets available, which allowed us to determine the baseline reproducibility (R^2^ = 0.55, RMSD = 1.52, **Figure S6A**). Applying the same MANOVA approach as above to map dependencies between protein positions and DNA positions, we found that the most significant associations are reproducible (**Figure S7A, B**). Next, we built a FamilyCode model and performed leave-one-out cross-validation of prediction of wild-type bHLH factors. Prediction performance for DNA position –1/+1 (R^2^ = 0.63, RMSD = 1.39) was better than that of the closest-paralog approach (R^2^ = 0.57, RMSD = 1.53), and similar as that of similarity regression (R^2^ = 0.62, RMSD = 1.45); see **Figure S6B-D** for details.

For a more rigorous validation of our ability to predict DNA binding specificity effects for unseen bHLH variants, we used PBM data for three unseen mutants of HLH-1 collected by De Masi et al. (59). Wild-type HLH-1 has Leu at bHLH domain residue position 13 and preferably binds to the CA<u>GC</u>TG E-box (**Figure S8A, B**); note that among the two available replicates for wild-type HLH-1 in CisBP, we chose the one with this same half-site preference (CAG). We predicted base preference shifts associated with the HLH-1 mutations L13R, L13T, and L13V (in the form of ΔΔΔG/RT values) from binding specificity models for wild-type bHLH factors alone using the same procedure used for HES2 and ASCL2 above, and compared these directly with empirical ΔΔΔG/RT values derived from the PBM data for the HLH-1 mutants (**Figure 4F, G**). Our prediction accuracy for motif position –1 was excellent (R^2^ = 0.98, RMSD = 0.19; see **Figure S8C**), and compared favorably to that of a state-of-the-art structure-based deep learning algorithm named DeepPBS (49) (R^2^ = 0.33, RMSD = 1.23; see **Figure S8D)**.

### Mapping DNA binding specificity determinants for homeodomains

To demonstrate the generalizability of our FamilyCode framework, we applied the same workflow to the homeodomain (HD) family of TFs. To obtain a more comprehensive training set, we used PyProBound to infer sequence-to-affinity models from all available HT-SELEX and PBM data for HDs (16, 33, 34, 51, 57). For HT-SELEX, a total of 259 unique HDs sourced from MotifCentral (22) had at least two replicates. For PBM, a total of 314 unique HDs are available in CisBP, of which 272 have a corresponding Uniprot entry. The final set of sequence-to-affinity models covered 414 unique HDs. We used 90 of these that were assayed using both SELEX and PBM to obtain a performance baseline (**Figure S9**).

Tetrahedral MANOVA analysis for this multi-platform compendium of HD binding models (**Figure 5A** and **Figure S7C, D**) identified several residue positions at which variation in amino-acid identity had been previously shown to impact DNA binding preference (27, 30). However, while most of these previous methods used structural information as part of their analysis, our MANOVA analysis is structure-agnostic.

**Figure 5:**
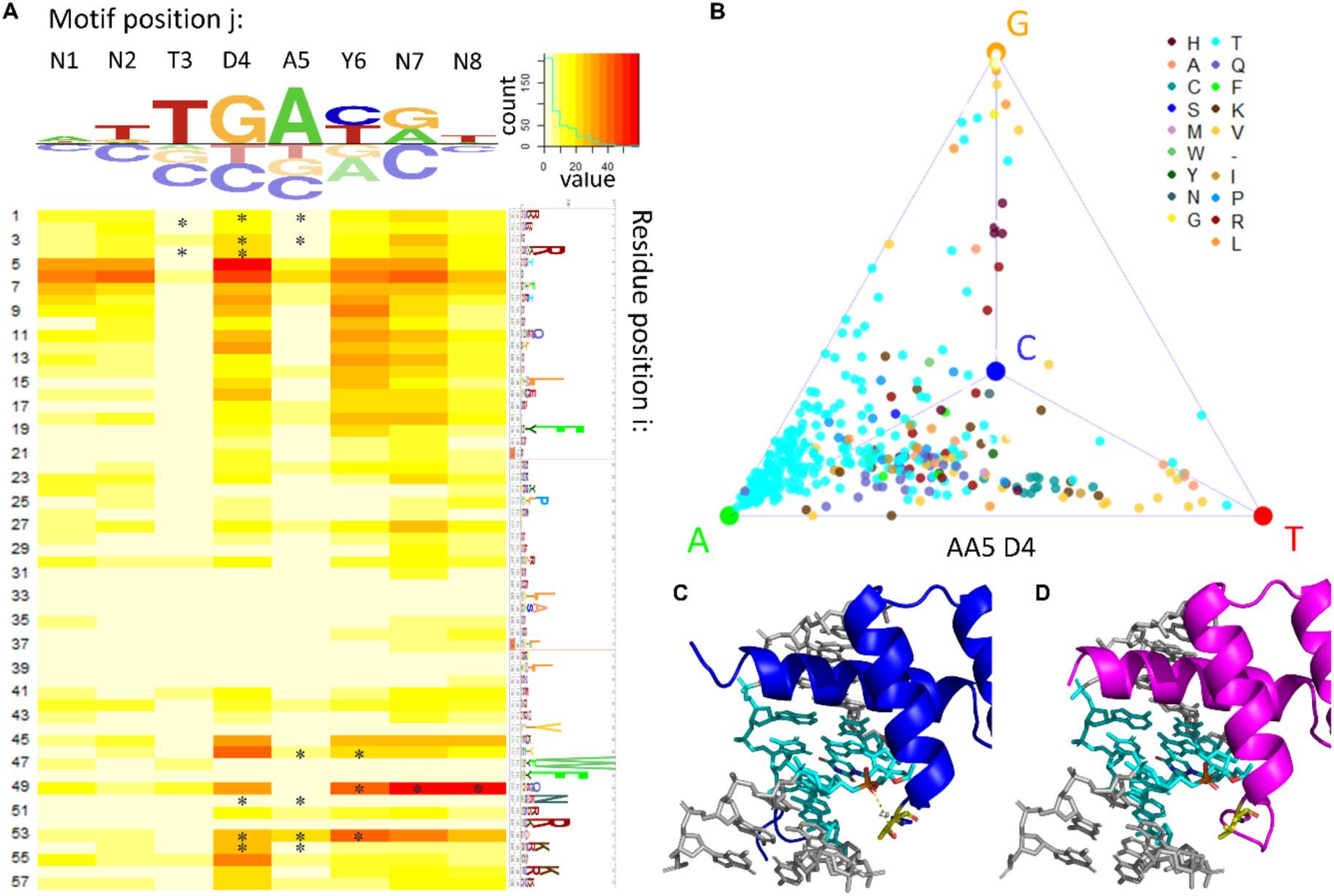
FamilyCode prediction for HD proteins using a model trained on HT-SELEX and PBM data. (**A**) Heatmap showing the p-value from a MANOVA test between amino-acid identity and position within the tetrahedron. Asterisks mark base contacting residues. (**B**) Tetrahedron representation of base preference at DNA position D4, with TFs colored according to amino-acid identity at residue position 5. (**C-D**) Structural inspection of the interaction between AA5 and D4 on Scr (**C**) and CDX1 (**D**), using PDB ID 2R5Z and 7Q3O.

Our finding that for HD positions 49 and 53 there is a strong association with base preference at DNA positions Y6 and N8 is consistent with previous findings (25, 27, 28, 48, 60), as is the finding that HD position 46 is associated with base preference at DNA position D4 (27, 28, 48). DNA-contacting HD positions 4 and 54 were not found to be associated with DNA binding specificity variation in our analysis due to their high conservation.

HD position 5 was previously identified as a highly associated residue in the prediction scheme by Christensen *et al.* (28), but was overlooked by structure-based approaches as residue 5 does not directly contact DNA (28). This position is highly associated with the base recognition at the D4 position of the motif, with Thr5 showing a consistent preference toward an A, while the hydrophobic residues Leu5, Ile5, and Val5 prefer either a G or a T (**Figure 5B**). For example, comparing the structures of Scr and CDX1, which have a Thr5 and Leu5 respectively, indicates that the side chain of AA5 can either form a hydrogen bond with the DNA backbone or not, depending on the amino acid identity (**Figure 5C, D**) (61). Christensen et al. and our method predict the same set of four HD positions as top determinants of DNA binding preference, which is satisfying because their data was from a different experiment platform (bacterial one-hybrid, or B1H). However, we note that our MANOVA approach is the only one that assesses the statistical significance of the identified positional associations.

### Predicting the effect of missense mutations in homeodomains on DNA binding specificity

A recent study (30) used PBM technology to profile how 92 different mutations in 30 HDs altered their DNA binding specificity, allowing us to directly and quantitatively assess our ability to predict the impact of mutations using our model as trained on wild-type HDs alone. To this end, we first used PyProBound to infer ΔΔG/RT models from the raw PBM intensities; this was done separately for each wild-type or mutant HD profiled in (30). In each case, we used the fraction of the variance in the raw PBM intensities explained by our model fit to select the highest-quality replicate (see **Supplemental Methods**). Subsequently, we used our tetrahedral representation to robustly estimate empirical ΔΔΔG/RT values for each HD mutant.

Figure S10A shows the empirical effect size the mutation in terms of the magnitude of the ΔΔΔG/RT range (difference between largest and smallest value) across all eight DNA positions, separately for each of the 92 mutants. There is a clear trend for mutants that were classified as having no effect on DNA binding by Kock et al. (30) to have a smaller impact according to our ΔΔΔG range metric (p=0.0024, Hypergeometric test with threshold of ΔΔΔG/RT = 1). An overview of the correlation in empirical ΔΔΔG/RT values between replicates is shown in **Supplemental Figure S10B**, and examples for specific mutants are shown in Figure 6B, C and **Supplemental Figure S11**.

**Figure 6:**
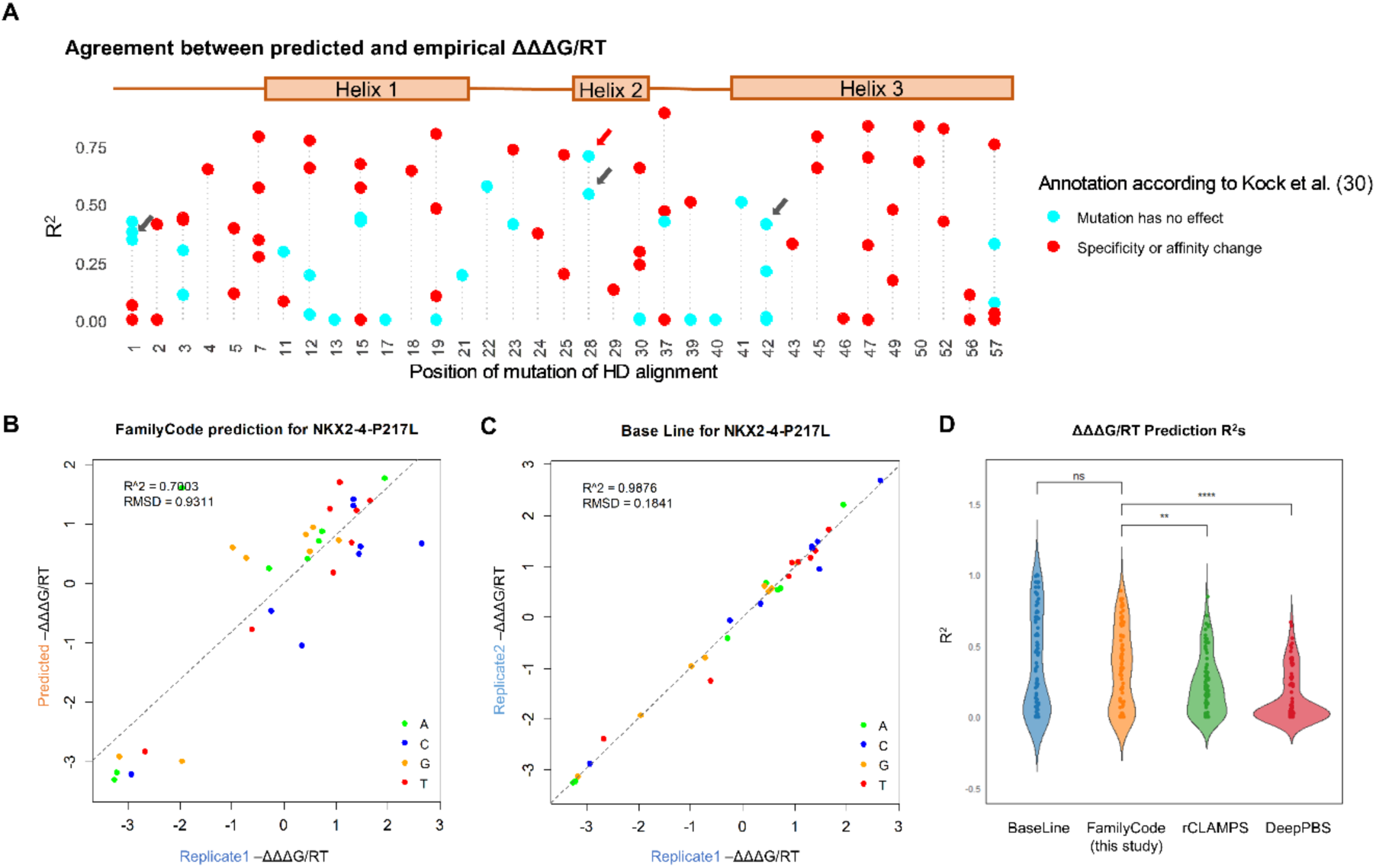
Validation of family-level predictions using disease-associated homeodomain mutants. (**A**) Predictive performance of FamilyCode as trained on wild-type homeodomains. Shown is the correlation (R^2^) between predicted and empirical ΔΔΔG/RT values for each of the 92 mutants profiled by Kock et al. (30). Arrows denote mutants for which we observed consistent effects on DNA binding specificity that have not previously been reported. (**B**) Replicate-to-replicate comparison of empirical ΔΔΔG/RT values for the P217L mutant of NKX2-4, indicated by the red arrow in panel A. (**C**) FamilyCode prediction vs. empirical ΔΔΔG/RT values for the same mutant. (**D**) Comparison of predictive performance of FamilyCode with rCLAMPS (48) and DeepPBS (49). Significance levels were computed using the paired Wilcoxon signed-rank test (ns: not significant; * p < 0.05; ** p < 0.01; *** p < 0.001; **** p < 0.0001).

Figure 6A shows how well the empirical ΔΔΔG/RT values for the 92 HD mutants can be predicted by FamilyCode when trained on wild-type HD data alone. **Figure S10** provides more information about the range and reproducibility of the empirical ΔΔΔG/RT values. Four mutants (indicated by grey arrows in Figure 6A and **Figure S10**) that were classified by Kock et al. as “no change” stood out in our own analysis as having a large empirical ΔΔΔG/RT range, excellent empirical ΔΔΔG/RT reproducibility, and good predictive performance of FamilyCode (Figure 6A, **Figure S10**, and **Figure S11**). This reanalysis of the same raw PBM data illustrates how the combination of PyProBound and our tetrahedron-based method for estimating ΔΔΔG/RT values allows for sensitive detection of mutant effects on DNA binding specificity.

The performance of FamilyCode (R^2^ for predicted v. empirical ΔΔΔG/RT values) across all 92 HD mutants is shown in orange in Figure 6D. It approaches the reproducibility in empirical ΔΔΔG/RT values inferred from two replicates for the same mutant HDs shown in blue. Our method has higher predictive power on the missense mutants than the state-of-the-art rCLAMPS model (48) for predicting HD specificity (p = 0.0057, rank sum test). The structure-based predictor DeepPBS (49) performed only modestly compared to both our method and rCLAMPS when applied to the same mutant HD validation set (Figure 6D). Note that to ensure a fair comparison, we used the same tetrahedral transformation to compute ΔΔΔG/RT values in all cases (see **Supplemental Methods**). It was previously shown that another structure-based prediction algorithm named ModCRE (62) is outperformed by rCLAMPS on the PWM prediction of TFs from the HD family. It is perhaps not surprising that family-specific prediction strategies such as FamilyCode and rCLAMPS outperform cross-family ones such as DeepPBS and ModCRE on family-specific prediction tasks.

## DISCUSSION

In this study, we introduced and validated a modeling framework for predicting the quantitative impact of missense mutations on the DNA binding specificity of transcription factors. Amino-acid substitutions can lead to loss of function related to a disruption of the TF’s ability to bind DNA, but our aim here was to predict more subtle quantitative effects on DNA binding preference, as these in turn might explain phenotypic effects associated with common or rare variants.

Each family of transcription factors is considered separately in our approach. While we focused on the bHLH and homeodomain families as a proof of concept, the approach is general, and could also be applied to other TF families that have a sufficiently larger number of members for which high-throughput DNA binding is available. We exploit the fact that amino-acid identity will vary at many positions in the protein alignment of wild-type transcription factors from a given family. Our family-level model estimates the average effect of amino-acid substitutions at each position using high-throughput *in vitro* binding data for each TF in the training set. We want to stress that our validation was focused on the effect of single-residue substitutions in the context of natural genetic variation. While our family-level models can also be used to make predictions for TFs that are many substitutions away from a wild-type reference, we expect that the accuracy of the predictions would be significantly lower in those cases. Therefore, our approach may be less suitable for the *de novo* engineering of novel TFs with prescribed DNA binding preferences.

An important determinant of the quality of our predictions is simply the total number of TFs in the training set, which was several dozen in our study. For larger families, a collection of human TF paralogs may suffice, but TFs from multiple species can be naturally combined in our approach. In fact, since the tetrahedral representation of base preference that we use as the basis for our family-level analysis is completely general, high-throughput binding data from multiple platforms (e.g., SELEX and PBM) can be seamlessly integrated in a single model. Moreover, as we have shown in previous work (22), high-throughput *in vivo* binding data such as ChIP-seq can yield models that are almost indistinguishable from those inferred from *in vitro* binding data as long as peak-agnostic analysis methods are used to infer the binding free energy coefficients. This implies that large-scale ChIP-seq data sets from projects such as ENCODE (63) might also be leveraged to increase the number of TFs in the training set for family-level modeling.

We want to stress that while the latest generation of structure prediction algorithms (50) are now capable of predicting the 3D structure of a protein-DNA complex from its primary sequence to a reasonable degree, this is still a vastly simpler task than predicting the energetic effect of base-pair or amino-acid substitutions at the protein-DNA interface. For the latter, there is currently no alternative to the combination of high-throughput experimental binding data combined with interpretable machine learning that we employed in this study. That said, there may still be a role for computational methods that predict the structure of the protein backbone: An implicit assumption in our family-level modeling is that the backbone geometry of the protein-DNA interface is fixed, and that the only thing that varies is the chemical identity of the base pairs and amino-acid sidechains. However, some TF families such as basic leucine zipper (bZIP) factors are capable of binding to DNA in alternative homodimer configurations (47, 64), and leveraging structure prediction methods to account for this in the context of family-level modeling may be a valuable extension of this approach.

Our approach is structure-agnostic in that it aims to directly predict TF *function* (i.e., base preference at a particular DNA position) from TF *sequence* (i.e., the amino acid identity at each position in the protein alignment). The limited structural data that we presented only serves to illustrate that our functional analysis yields result that make mechanistic sense. The protein sequence alignment only serves to link corresponding residue positions across TFs. We do not use knowledge about the evolutionary relationships between the various TFs in the alignment, although these do lead to correlations between the binary amino-acid indicators that we use as independent variables in our family model, which is one of the reasons why we use forward feature selection. It is conceivable that foundation models built from large sets of homologous protein sequences (65) can be leveraged to create a more informative mapping of individual TF protein sequences to a vector of continuous feature weights, thereby increasing the predictive performance of our family-level models. Finally, it has been demonstrated empirically that an equivalent missense substitution can have different effects on DNA binding in different HD mutants (30), and multi-domain zinc finger (ZF) transcription factors are known to exhibit strong dependencies between adjacent ZF domains (66, 67). It would be interesting to leverage the tetrahedron representation to account for such dependence on protein sequence context when predicting mutation effects based on wild-type training data.

## LIST OF SUPPLEMENTAL DATA

**Supplemental Data S1**: DNA recognition models for the 52 bHLH factors analyzed in this study.

**Supplemental Data S2:** Interactive 3D representations of tetrahedrons. Open HTML files in browser to view and manipulate. Related to Figure 2A-C.

**Supplemental Data S3:** Empirical cumulative distribution of tetrahedral position along each principal component direction for various amino-acid positions. Related to Figure 4B.

**Supplemental Data S4:** Statistical significance of ANOVA test of PCs at DNA position –1/+1, –2/+2 and –3/+3.

**Supplemental Data S5:** JSON configuration file used for all ProBound analyses.

All supplementary data files are available for download at https://bussemakerlab.org/papers/FamilyCode/.

## ACKNOWLEDGEMENTS

The research reported in this publication was supported by NIMH award R01MH106842 and NHGRI award R01HG003008 to H.J.B. and NIGMS award R35GM118336 to R.S.M. We thank members of the Bussemaker and Mann labs for useful discussions. We also thank Todd R. Riley for useful discussion during the early stages of this project.

## AUTHOR CONTRIBUTIONS

MPG and HJB conceived of the project; SL, MPG, and HJB developed computational methods; SL, MPG, and CL wrote computer code for family-level modeling; LANM implemented the PBM extension of PyProBound; SL performed all computational analyses under the supervision of HJB; XL provided guidance for structural analyses; SL performed all wet lab experiments with input from WJG and under the supervision of RSM; SL and HJB wrote the paper with significant input from RSM; all authors edited and approved the paper.

## DATA AND SOFTWARE AVAILABILITY

All computer code developed for and used in this study, including scripts to generate all figure panels from scratch, is available at http://github.com/BussemakerLab/FamilyCode. Raw sequencing data for the SELEX-seq assays we performed in this study was submitted to the NCBI short read archive (BioProject identifier PRJNA1244358; available to reviewers via <u>this link</u>).

---

**Figure S1:**
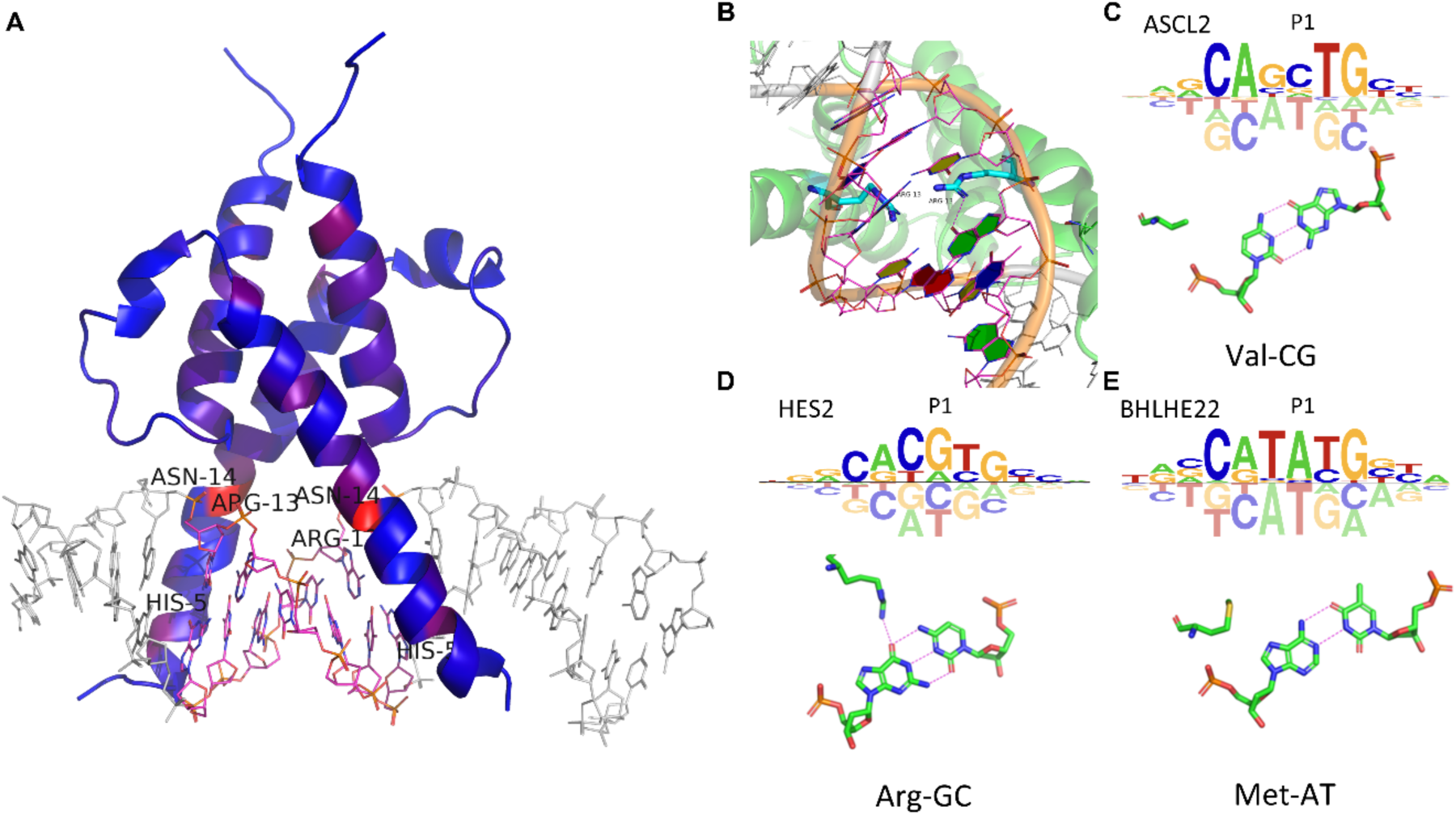
Structural analysis of bHLH DNA-binding residues. (**A**) Statistical significance at position –1 shown in the context of the co-crystal structure for Pho4 (red denotes significant p-values). (**B**) Detailed view of the interaction between Arg13 and a GC base pair at motif position +1. The Arg13 sidechain is colored in cyan. The E-box is shown in magenta using DNA blocks. (**C**-**E**) Additional examples of interactions between amino-acids and base pairs at residue position 13 and DNA position +1. Structural images generated using 3DNA/DSSR (x3dna.org)(68). Hydrogen bonds are shown as pink dashed lines.

**Figure S2:**
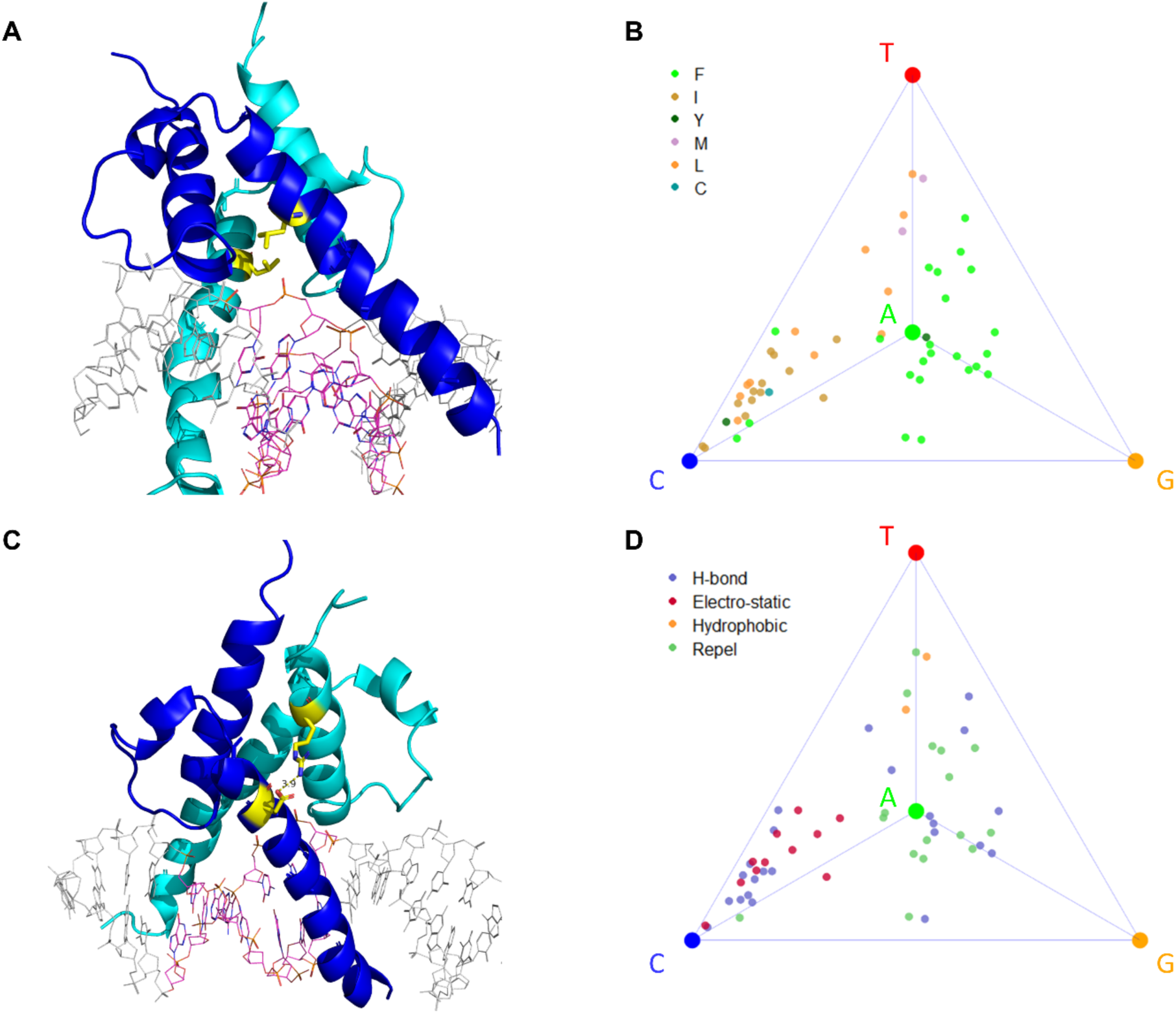
Associations found for residue positions 20 and 50 in the bHLH family analysis. (**A**) Structure of bHLH protein PHO4 with residue position 20 highlighted in yellow, and blue and cyan indicate the two bHLH monomer subunits. (**B**) Tetrahedron representation for DNA position –1, with coloring according to the amino acid identity at residue position 20. (**C**) Same as panel A, but for residue positions 22 and 50. (**D**) Tetrahedron representation of DNA position –1 with coloring according to the type of contact between residue positions 22 and 50.

**Figure S3:**
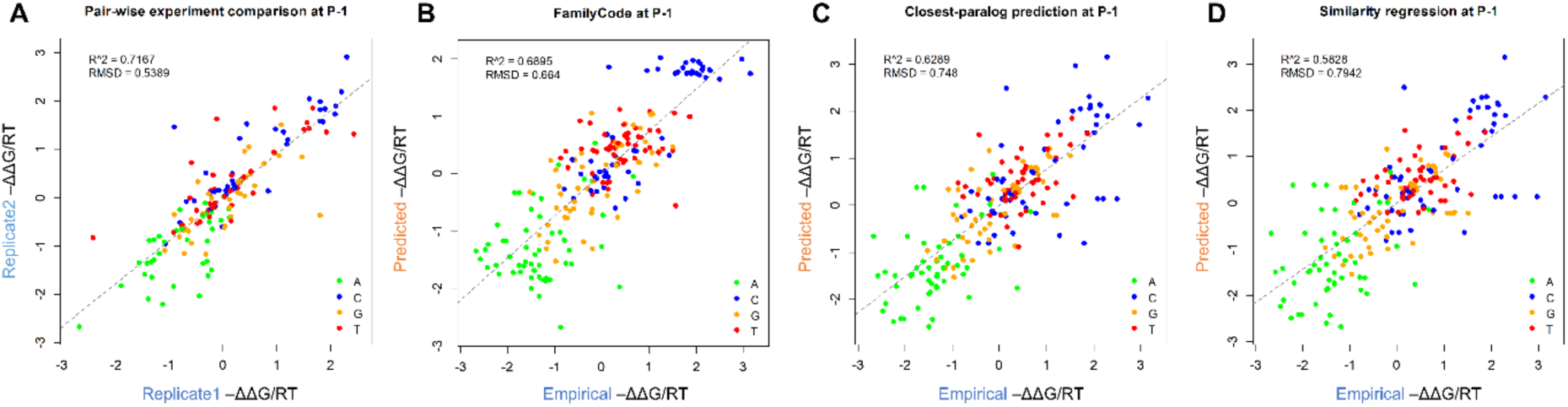
Cross-validation on wild-type bHLH binding motifs generated from HT-SELEX data. (**A**) Reproducibility between HT-SELEX replicates. (**B-D**) Comparison between empirically determined binding energies and predictions made using our family-code (**B**), the closest-paralog method (**C**), or similarity regression (**D**).

**Figure S4:**
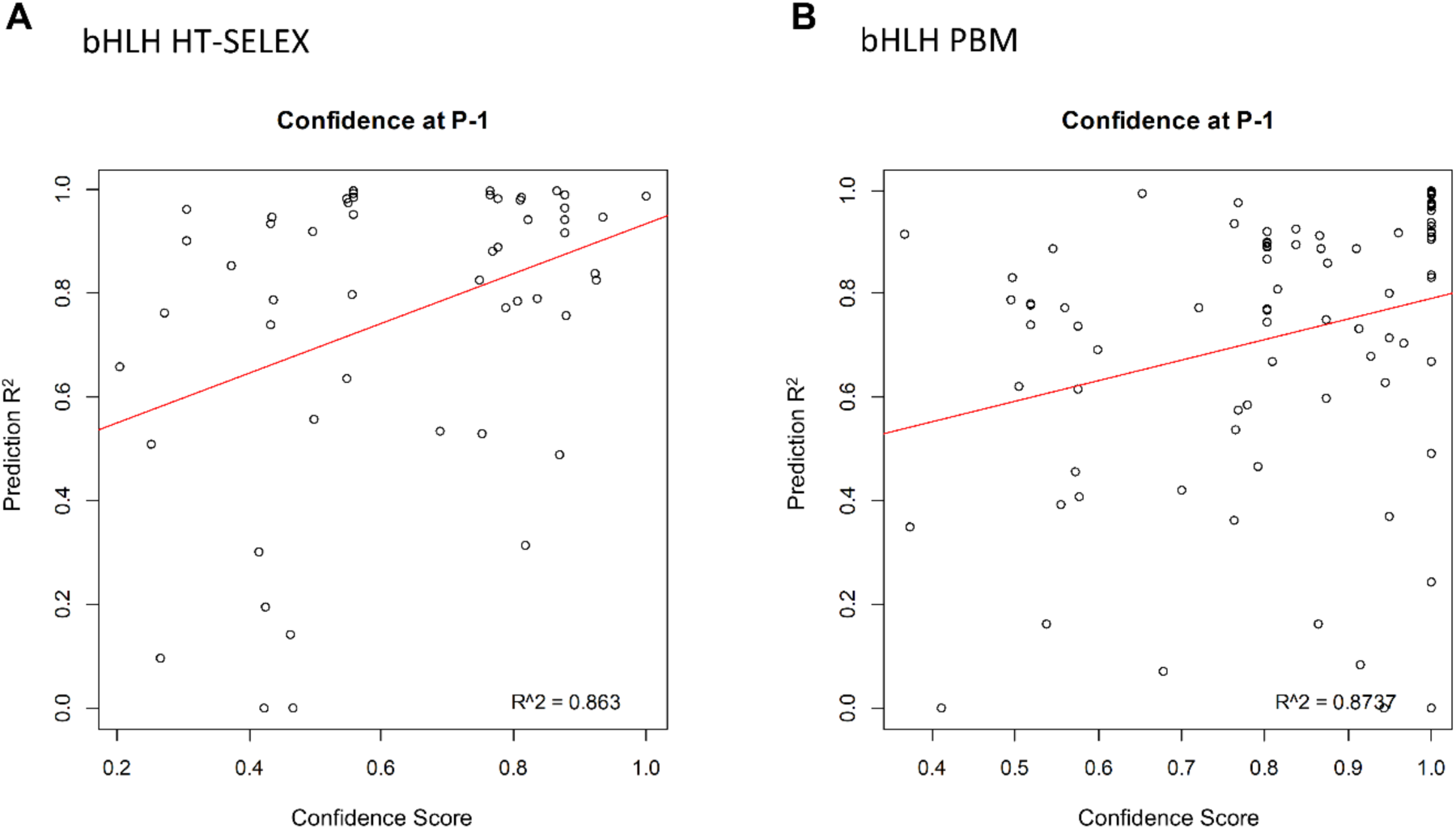
Correlation between prediction accuracy and confidence score. For bHLH models derived from HT-SELEX (**A**) and PBM (**B**) data.

**Figure S5:**
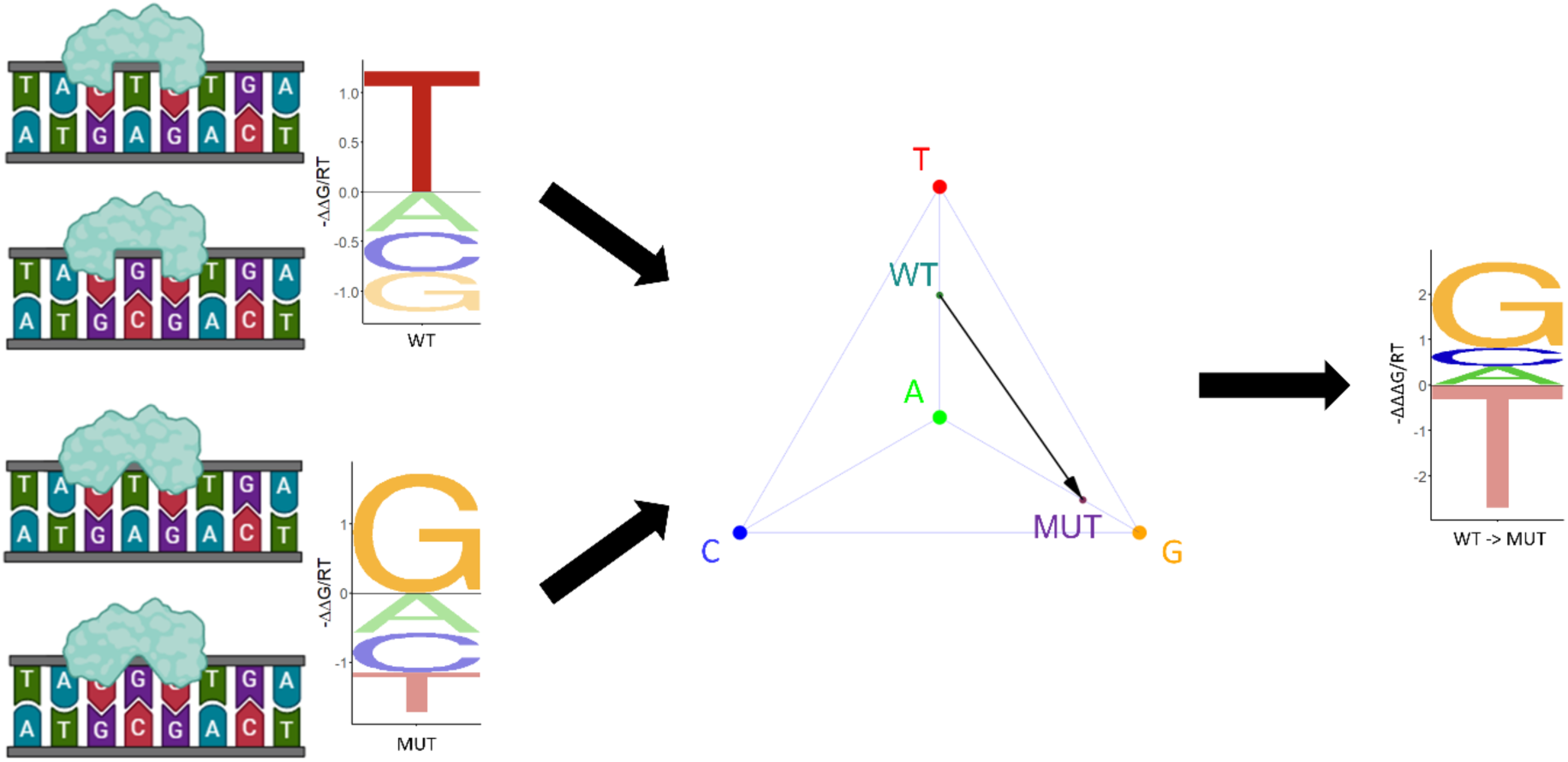
Illustration of our tetrahedron-based procedure for calculating ΔΔΔG/RT values.

**Figure S6:**
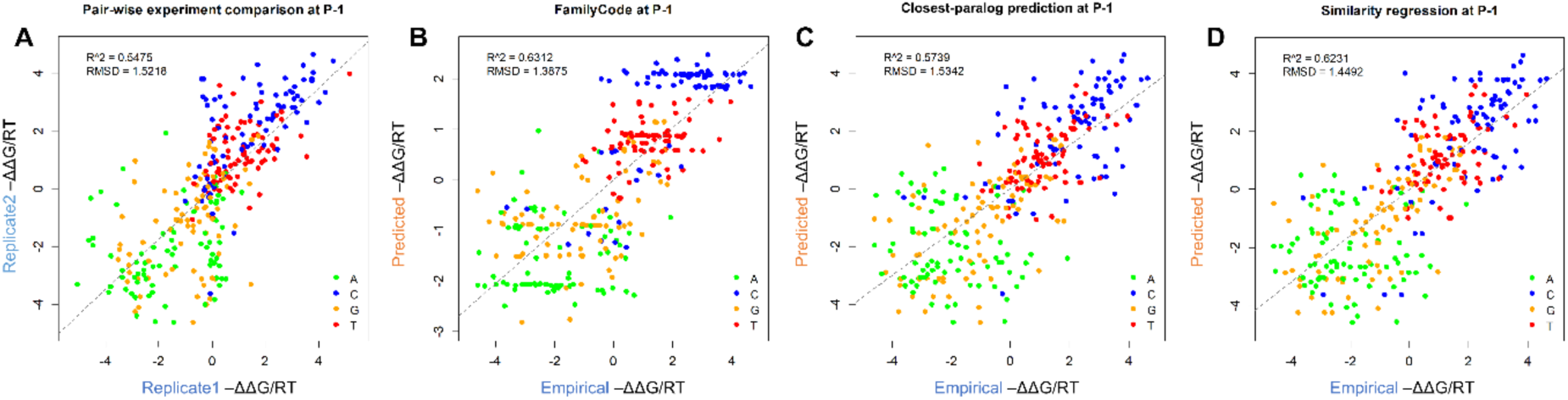
Cross-validation on wild-type bHLH binding motifs generated from PBM data. (**A**) Reproducibility between PBM replicates. (**B-D**) Comparison between empirically determined binding energies and predictions made using our family-code (**B**), the closest-paralog method (**C**), or similarity regression (**D**).

**Figure S7:**
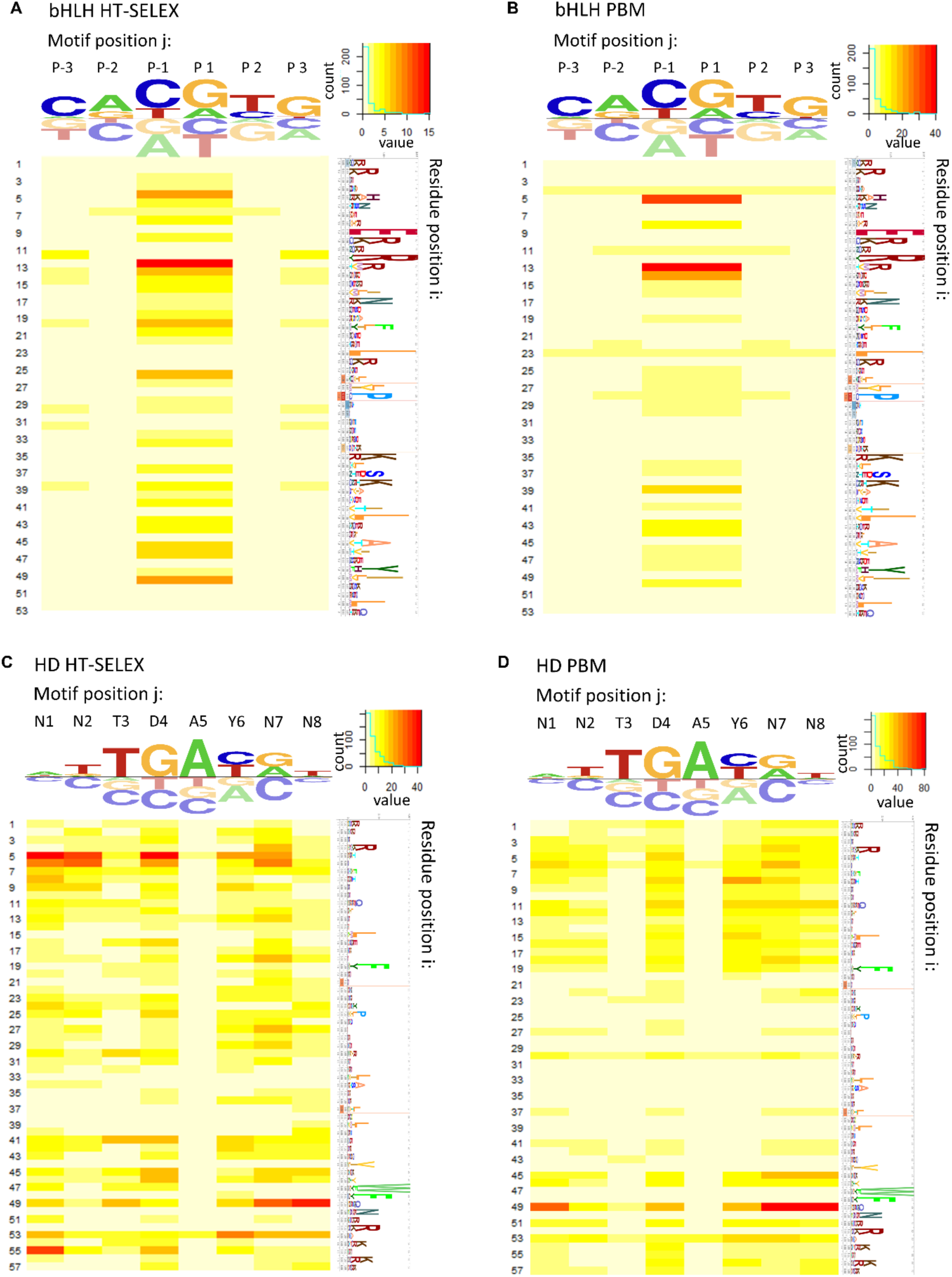
Positional association maps. Shown are results for the bHLH (**A**, **B**) and HD (**C**, **D**) families generated using HT-SELEX (**A**, **C**) and PBM (**B**, **D**) data.

**Figure S8:**
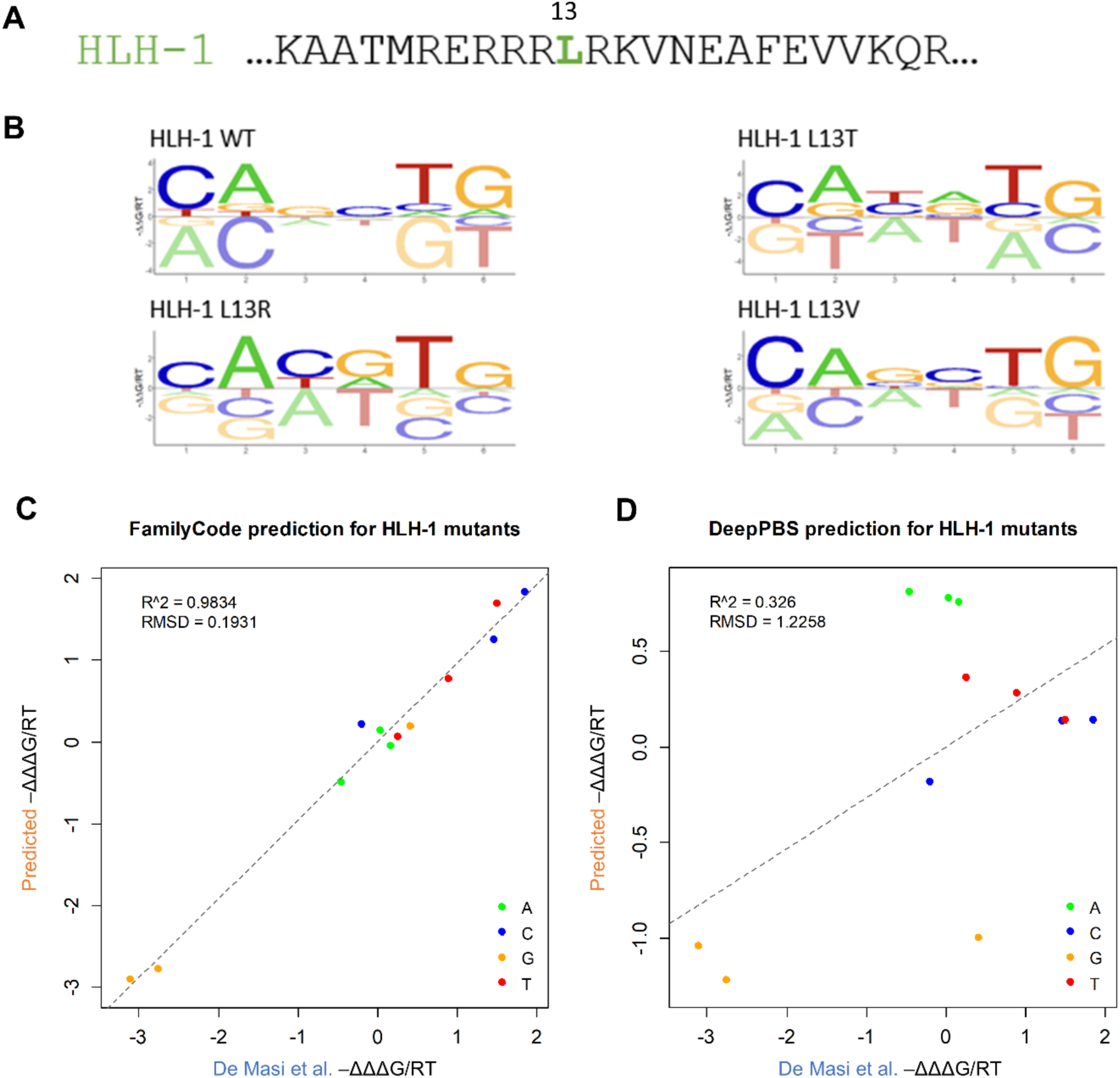
Family code predictions for mutant HLH-1. (**A**) Wild-type sequence of *C. elegans* factor HLH-1, with mutated position 13 indicated. (**B**) Energy logos for wild-type and mutant HLH-1 factors inferred from PBM data using PyProBound. (**C**) Predicted versus empirically values of ΔΔΔG for the three HLH-1 mutants.

**Figure S9:**
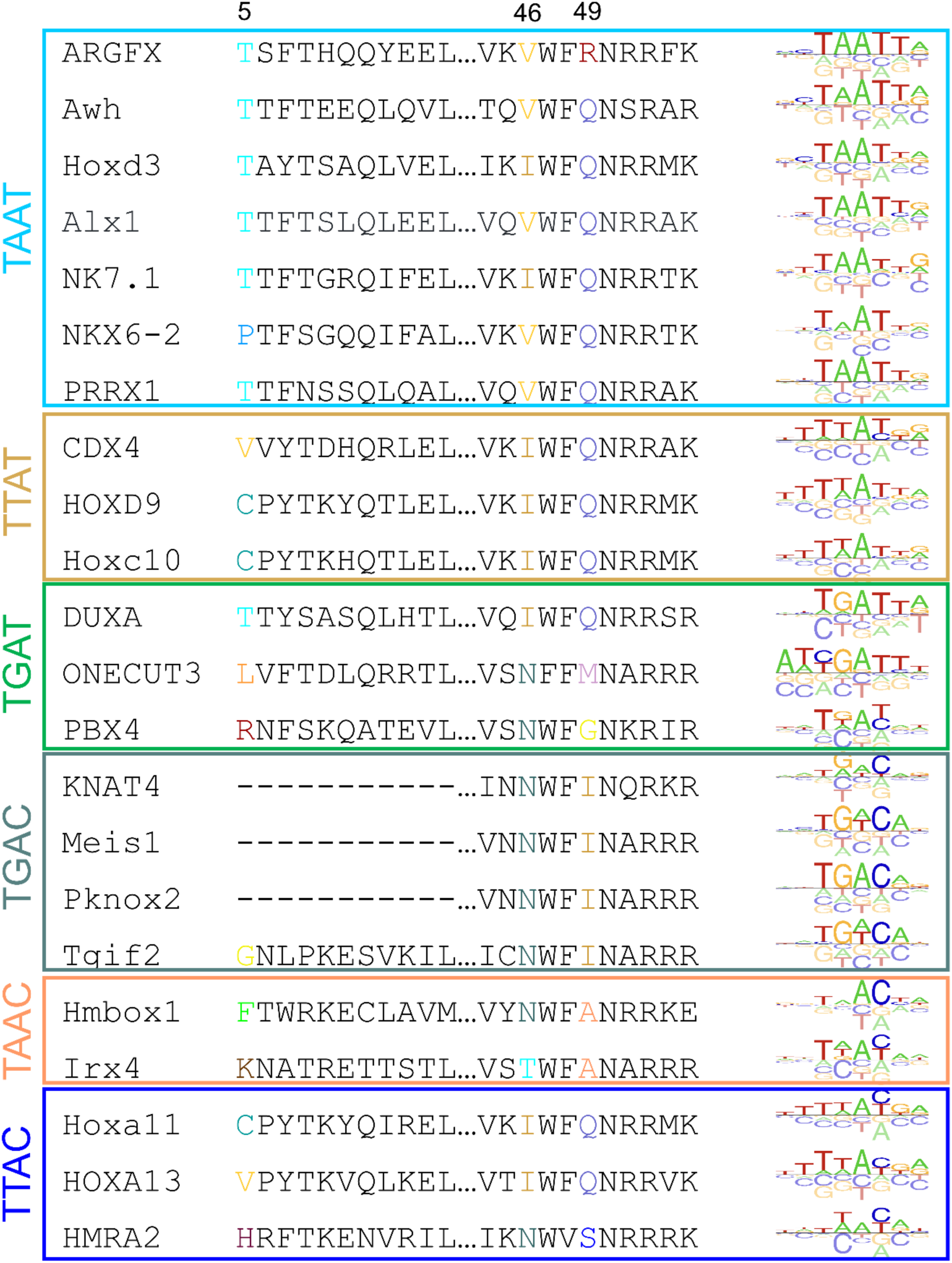
Variation in DNA binding preference within the Homeodomain (HD) family of transcription factors. Aligned protein sequences and DNA binding energy logos for a representative set of HD factors, grouped by preferred homeobox core.

**Figure S10:**
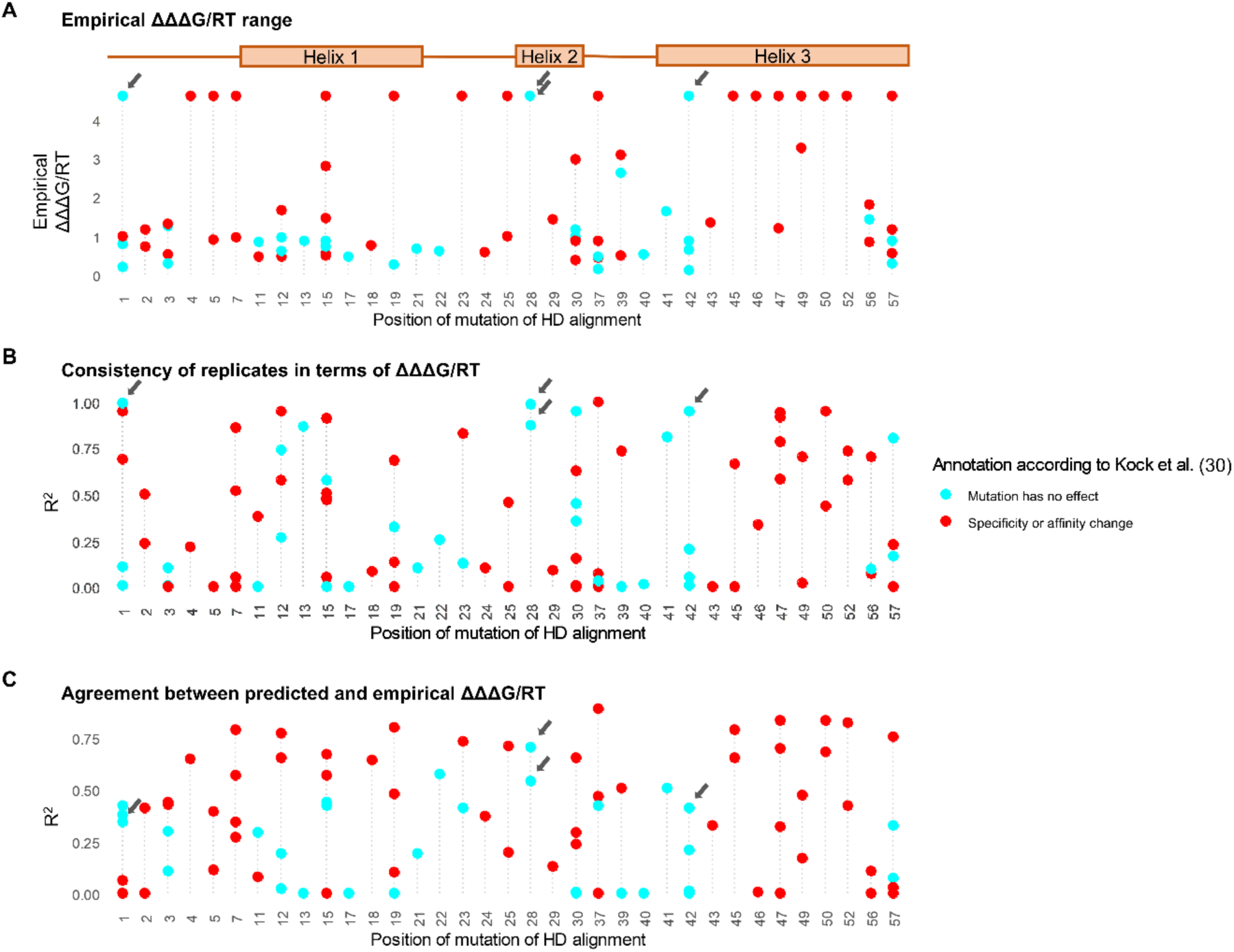
Reanalysis of PBM data for a set of 92 mutant homeodomains. (A) ΔΔΔG/RT range of empirical motif model plotted along mutation position. Wilcoxon signed-rank test of ΔΔΔG/RT ranges between reported change and unchanged sample resulted in p = 0.0006357. (B) Empirical R^2^ between two replicates of each mutant. (C) Prediction R^2^ between family code prediction ΔΔΔG model and replicate 1 of empirical model. Grey arrows annotate mutations that have outstanding specificity determining effects that are not discovered in the original Kock et al. study, and are shown in detail in **figure S11**.

**Figure S11:**
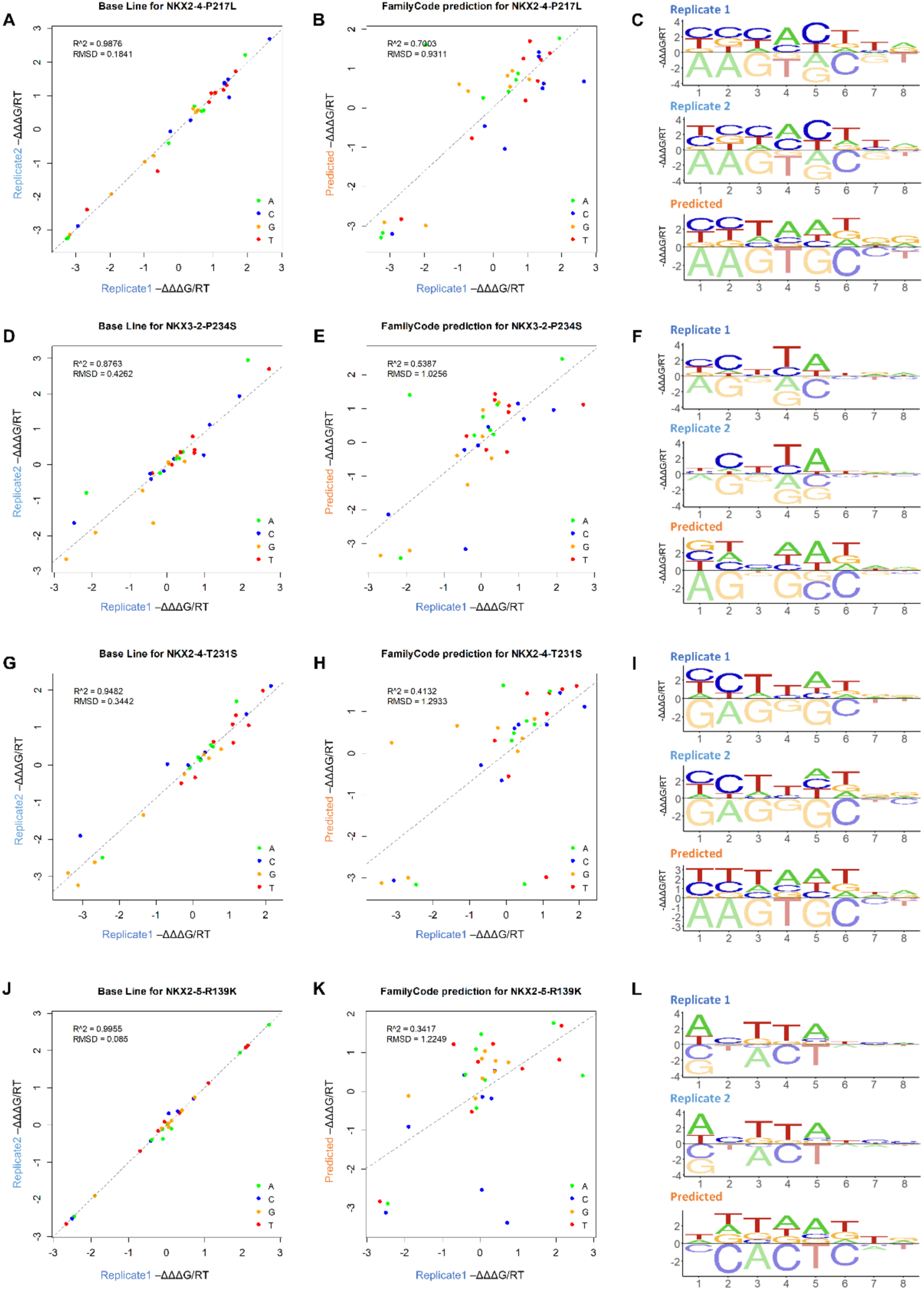
(**A**) Family code prediction vs. replicate 1 for NKX2.4-P217L (P28L). (**B**) Replicate 1 vs. replicate 2 for NKX2.4-P217L (P28L). (**C**) Empirical and predicted ΔΔΔG models for NKX3.2-P234S (P28S). (**D**) Family code prediction vs. replicate 1 for NKX3.2-P234S (P28S). (**E**) Replicate 1 vs. replicate 2 for NKX3.2-P234S (P28S). (**F**) Empirical and predicted ΔΔΔG models for NKX3.2-P234S (P28S). (**G**) Family code prediction vs. replicate 1 for NKX2.4-T231S (T42S). (**H**) Replicate 1 vs. replicate 2 for NKX2.4-T231S (T42S). (**I**) Empirical and predicted ΔΔΔG models for NKX2.4-T231S (T42S). (**J**) Family code prediction vs. replicate 1 for NKX2.5-R139K (R1K). (**K**) Replicate 1 vs. replicate 2 for NKX2.5-R139K (R1K). (**L**) Empirical and predicted ΔΔΔG models for NKX2.5-R139K (R1K).

**Figure S12:**
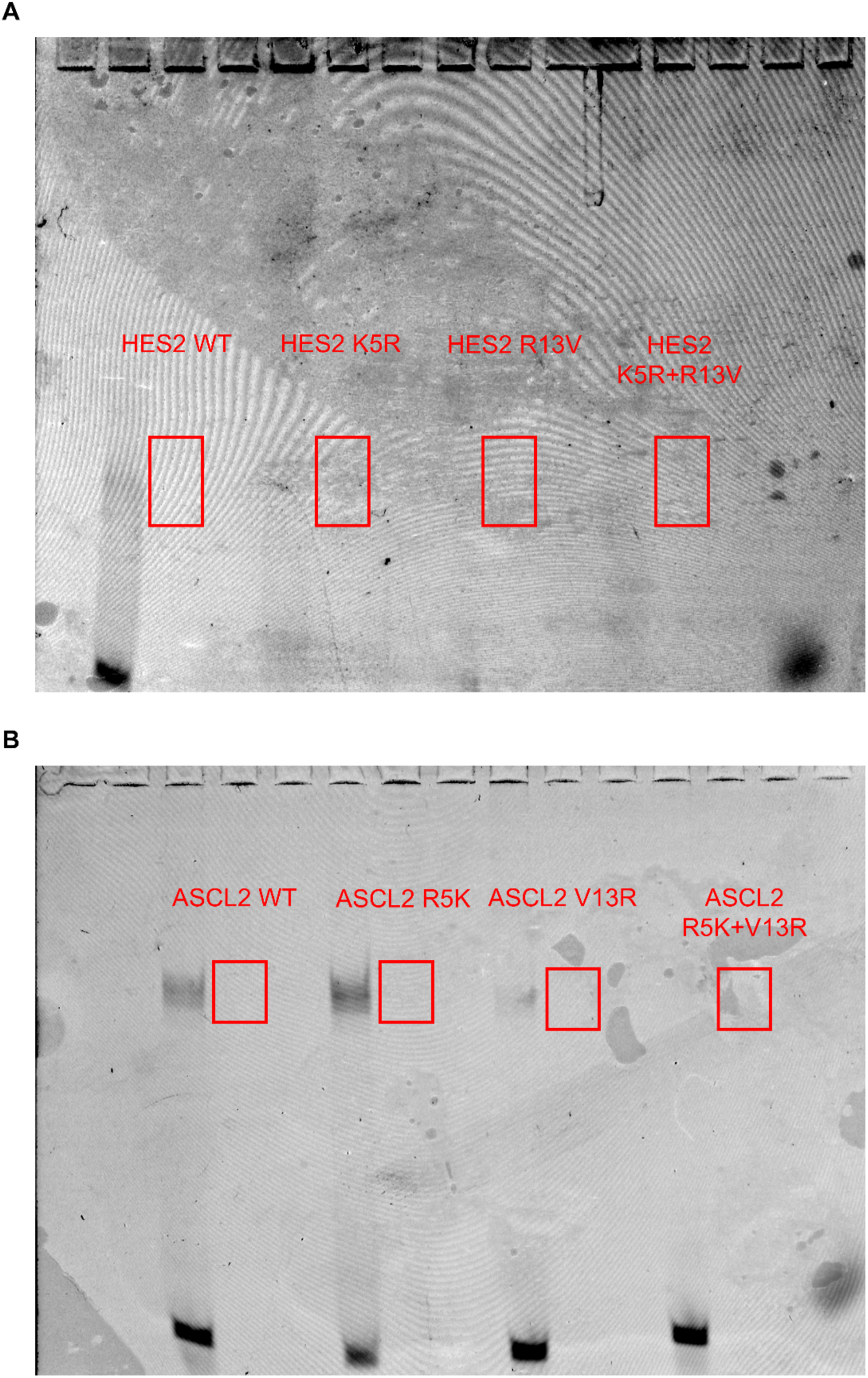
Raw EMSA gels. Shown are the bands we cut out to perform SELEX-seq for wild-type and mutant HES2 (**A**) and ASCL2 (**B**).

